# Riluzole suppresses growth and enhances response to endocrine therapy in ER+ breast cancer

**DOI:** 10.1101/2020.07.30.227561

**Authors:** Ayodeji O. Olukoya, Hillary Stires, Shaymaa Bahnassy, Sonali Persaud, Yanira Guerra, Suman Ranjit, Shihong Ma, M. Idalia Cruz, Carlos Benitez, Aaron M. Rozeboom, Hannah Ceuleers, Deborah L. Berry, Britta M. Jacobsen, Ganesh V. Raj, Rebecca B. Riggins

## Abstract

**Background:** Resistance to endocrine therapy in estrogen receptor-positive (ER+) breast cancer remains a significant clinical problem. Riluzole is FDA-approved for the treatment of amyotrophic lateral sclerosis. A benzothiazole-based glutamate release inhibitor with several context-dependent mechanism(s) of action, Riluzole has shown anti-tumor activity in multiple malignancies, including melanoma, glioblastoma, and breast cancer. We previously reported that the acquisition of Tamoxifen resistance in a cellular model of invasive lobular breast cancer is accompanied by the upregulation of GRM mRNA expression and growth inhibition by Riluzole.

**Methods:** We tested the ability of Riluzole to reduce cell growth, alone and in combination with endocrine therapy, in a diverse set of ER+ invasive ductal and lobular breast cancer-derived cell lines, primary breast tumor explant cultures, and the estrogen-independent, *ESR1*-mutated invasive lobular breast cancer patient-derived xenograft model HCI-013EI.

**Results:** Single-agent Riluzole suppressed the growth of ER+ invasive ductal and lobular breast cancer cell lines *in vitro*, inducing a histologic subtype-associated cell cycle arrest (G0-G1 for ductal, G2-M for lobular). Riluzole induced apoptosis and ferroptosis and reduced phosphorylation of multiple pro-survival signaling molecules, including Akt/mTOR, CREB, and Src/Fak family kinases. Riluzole, in combination with either Fulvestrant or 4-hydroxytamoxifen, additively suppressed ER+ breast cancer cell growth *in vitro*. Single-agent Riluzole significantly inhibited HCI-013EI patient-derived xenograft growth *in vivo*, and the combination of Riluzole plus Fulvestrant significantly reduced proliferation in primary breast tumor explant cultures.

**Conclusions:** Riluzole, alone or combined with endocrine therapy, may offer therapeutic benefits in diverse ER+ breast cancers, including lobular breast cancer.

## Background

Estrogen receptor-positive (ER+) breast cancer is the most commonly diagnosed cancer among women in the United States [1]. Endocrine therapies ranging from selective estrogen receptor modulators and downregulators (SERMs, SERDs) to aromatase inhibitors are the backbone of our current standard of care for the clinical management of ER+ breast cancers [2]. Although these treatments have significantly improved disease-free and overall survival for individuals with ER+ breast cancer, endocrine resistance remains a persistent, multifactorial problem [3]. Current efforts aim to address this problem through treatment with endocrine agents combined with other molecularly targeted therapies.

To further complicate these efforts, ER+ breast cancer is not a single disease. Invasive lobular breast cancer (ILC) is a distinct histologic subtype of breast cancer that is overwhelmingly ER+ yet has distinct genomic, transcriptomic, and proteomic features [4–6]. These distinctions have important implications for endocrine therapy response. Also, when compared to the more common invasive ductal breast cancer (IDC, invasive mammary carcinoma of no special type), ILC carries a greater risk for late recurrence (evident > 6 years after initial diagnosis) [7,8] and responds less to the SERM Tamoxifen [9,10] and potentially the steroidal aromatase inhibitor exemestane [11]. Additionally, models of ILC are less responsive to the second-generation SERD AZD9496 than Fulvestrant, while these drugs are equipotent in preclinical models of IDC [12].

Our group [13,14] and others [12,15–20] have identified a number of potential mechanisms that contribute to endocrine therapy resistance in ILC. We recently identified the upregulation of multiple metabotropic glutamate receptors (mGluRs, GRMs) in Tamoxifen-resistant ILC cells [14]. This, coupled with other studies that directly or indirectly implicate altered amino acid metabolism and signaling in ILC pre-clinical models [21] and clinical disease [22,23], led us to consider whether glutamate signaling is functionally relevant to endocrine resistance in endocrine-resistant ILC. Initially reported in melanoma [24,25] and now other malignancies (e.g.[26]), pro-tumorigenic signaling through GRMs can be inhibited by Riluzole, an oral benzothiazole-based glutamate release inhibitor that is FDA-approved for the treatment of amyotrophic lateral sclerosis (ALS). Riluzole’s proposed mechanism of action within the central nervous system in ALS and in melanoma is that blocking glutamate release into the extracellular space starves GRMs of their glutamate ligand, thus functionally inhibiting them. This inhibition of the GRMs ultimately reduces glutamate excitotoxicity and inhibits tumor cell growth. In triple-negative breast cancer (TNBC), Riluzole’s action may not depend on GRMs [27,28], although Riluzole exerts anti-tumor effects [29–31]. Despite the potential of repurposing Riluzole in ER+ breast cancer, especially ILC, this approach has not been a major focus to date. Therefore, this study aims to more broadly test the efficacy of Riluzole, alone and in combination with multiple endocrine therapies, in a diverse set of ER+ *in vitro* and *in vivo* models enriched for ILC.

## Methods

### Cell Culture and Reagents

Several cell lines were cultured, maintained, and used in this study. These cell line models include; ER+ ILC cell lines (SUM44, LCCTam, MDA-MB-134VI (MM134) and MDA-MB-134VI long-term estrogen-deprived (LTED; MM134 LTED), and BCK4), ER+ IDC cell lines (MCF7 and LCC9), and the ER- negative (ER-) non-transformed mammary epithelial cell line MCF10A as a control. MCF7 and LCC9; SUM44 and LCCTam; MM134 and MM134 LTED all represent pairs of parental and resistant cell lines, respectively. SUM44 and LCCTam cells were maintained under serum-free conditions in improved minimal essential media (IMEM, #A1048901, ThermoFisher, Grand Island, NY) supplemented with insulin, hydrocortisone, and other supplements as previously described [13], with the addition of 500 nM 4-hydroxytamoxifen (#H7904, Sigma Aldrich, St. Louis, MO) to LCCTam cells. For selected experiments, SUM44 and LCCTam cells were maintained under serum-free conditions in a base media of Ham’s F12 (#11765062, ThermoFisher) supplemented as above. MM134 cells and MM134 LTED) were maintained in IMEM supplemented with 10% fetal bovine serum (FBS) and phenol red-free IMEM (#A1048801, ThermoFisher) supplemented with 10% charcoal-cleared serum (CCS), respectively. BCK4 cells were maintained in IMEM supplemented with 10% FBS, insulin, and nonessential amino acids, as previously described [32]. MCF7 and LCC9 cells were maintained in IMEM supplemented with 5% FBS and phenol red-free IMEM supplemented with 5% CCS, respectively. MM134 and MCF7 cells were short-term hormone-deprived for selected experiments by culturing in phenol red-free IMEM supplemented with 5% CCS for 72 hours. The immortalized mammary epithelial cell line MCF10A was maintained as previously described [33]. All cell lines were authenticated by short tandem repeat (STR) profiling and regularly tested to ensure they remained free of *Mycoplasma spp*. contamination. Unless otherwise noted, general cell culture supplements and reagents were purchased from either ThermoFisher or Sigma Aldrich. Fulvestrant and Riluzole were purchased from Sigma Aldrich, Tocris Bio-Techne (Minneapolis, MN), or Selleckchem (Houston, TX). Ferrostatin-1 was purchased from Selleckchem.

### Cell Proliferation Assays

On Day 0, cells were seeded in 96-well plates at the following densities: 1,000 cells/well (MCF7); 2,000 cells/well (LCC9, MCF10A); 10,000 cells/well (SUM44, LCCTam, MM134, MM134 LTED); 15,000 cells/well (BCK4). Forty-eight hours later, on Day 2, cells were treated with the indicated concentration of compound(s) or solvent control (DMSO) for an additional 7 or 8 days, with media/compound(s) replaced on Day 5 or 6. Plates were then stained with crystal violet, dried, rehydrated, and read as previously described in [14]. Data are presented as mean ± standard error of the mean (SEM, Riluzole growth curves), or median with upper/lower quartiles (effect of 10 μM Riluzole on cell line pairs) of % growth (vehicle = 100%) for 5-6 technical replicates and are representative of 2-4 independent biological assays. For assays of Riluzole in combination with Fulvestrant or Tamoxifen for all cell lines except BCK4, data are processed as the mean % growth for 5-6 technical replicates and represent 2-4 independent biological assays. A representative single technical replicate’s mean % growth data was then used to create a combination matrix used in SynergyFinder, and the results were presented as 2D surface plots. SynergyFinder uses predictive models such as highest single agent (HSA), Bliss, and Zero interaction potency (ZIP) to quantify the degree of combination synergy or antagonism and outputs a synergy score. When interpreting SynergyFinder scoring, a synergy score less than −10 shows antagonistic drug interaction, a score between −10 and 10 shows an additive drug interaction and a score greater than 10 shows a synergistic drug interaction. For the assay of Riluzole in combination with Fulvestrant in BCK4 cells, data are presented as median with upper/lower quartiles of % growth for 5-6 technical replicates and represent 2-4 independent biological assays.

### Cell Cycle Assays

On Day 0, cells were seeded in 6-well plates at the following densities: 150,000 cells/well (MCF7, LCC9); 300,000 cells/well (SUM44, LCCTam, MM134, MM134 LTED, BCK4, MCF10A). Forty-eight hours later, on Day 2, cells were treated with 10 μM Riluzole or DMSO control for the additional indicated times before collection, fixation, staining, and cell cycle analysis by flow cytometry as described in [34]. Data are presented as mean ± standard deviation (SD) for 3-4 independent biological assays.

### Annexin V Apoptosis Assays

On Day 0, cells were seeded in a 6-well plate at 300,000 cells/well (SUM44, LCCTam). Forty-eight hours later, on Day 2, cells were treated with either 10 μM Riluzole or DMSO solvent as a control for another 48 hours. On Day 4, cells were collected and stained with 4μL of propidium iodide (PI) and 4μL of annexin V conjugated with fluorescein isothiocyanate (FITC) in 100μL of 1X binding buffer. Control cells were either left unstained or stained with either PI or annexin V dye conjugated with FITC, as described in [35]. Live (PI^-^, annexin V^-^), early apoptotic (PI^-^, annexin V^+^), late apoptotic(PI^+^, annexin V^+^), and necrotic cells (PI^+^, annexin V^-^) were quantified by flow cytometry. The PI/Annexin V-FITC apoptosis detection kit was purchased from BioLegend (#640914, San Diego, CA, USA). Data are presented as mean ± SD for 3 (SUM44) or 4 (LCCTam) independent biological assays.

### Human Phospho-Kinase Proteome Profiler™ Array

On Day 0, cells were seeded in a 6-well plate at 500,000 cells/well (SUM44, LCCTam). Forty-eight hours later, on Day 2, cells were treated with either 10 μM Riluzole or DMSO solvent as a control for 48 hours. On Day 4, cells were collected in 100 μL lysis buffer/well before determining total protein concentration by bicinchoninic acid (BCA) assay (#23225, ThermoFisher). According to the manufacturer’s instructions, five hundred micrograms of whole cell lysate were then assayed using the Human Phospho-Kinase Proteome ProfilerTM Array (#ARY003B, Bio-Techne). Array membranes were visualized using chemiluminescence detected by HyBlot CL autoradiography film (#E3018, Thomas Scientific, Swedesboro, NJ), then films were scanned and analyzed using FIJI [36]. A ratio of background-corrected intensity values for targets (phospho-kinase spots) to references (control spots) was created for each condition (DMSO and Riluzole) within each cell line. Data are presented as the mean of the Riluzole: DMSO ratio for two technical replicates from a single experiment for each cell line. Gene symbols corresponding to the kinases showing decreased phosphorylation in response to Riluzole in LCCTam cells were analyzed using SRPlot (http://www.bioinformatics.com.cn/srplot) to identify top functional enrichments.

### Western blot

SUM44 and LCCTam cells were seeded in 6-well plastic tissue culture plates at 250,000 cells/well (FAK and YES blots) or 300,000-600,000 cells/well (4-HNE and MDA blots) 48 h before treatment. The cells were treated for the times indicated in the figure legend. For the FAK and YES blots, the cells were treated with the control (DMSO) or drug (10μM Riluzole). In the case of the MDA and 4-HNE blots, the cells were treated with control (DMSO), Riluzole (10 μM), or a combination of Riluzole and Ferrostatin-1(10 μM). After treatment, cells were lysed in radioimmunoprecipitation assay buffer (RIPA - 150 mM NaCl, 50 mM Tris pH 7.5, 1% Igepal CA-630, and 0.5% sodium deoxycholate) supplemented with Pierce™ protease and Phosphatase inhibitor mini-tablets (Thermo Scientific). Protein lysates, extracted following centrifugation of the lysed cells, were mixed in a 3:1 with sample buffer (NuPAGE™ LDS Sample Buffer (4X) + 2-Mercaptoethanol in 2:1) and loaded onto a precast Gel (NuPAGE™ 4-12% Bis-Tris Gel, Invitrogen). Proteins were transferred to nitrocellulose membranes, blocked in 5% nonfat dry milk in Tris Buffered Saline and Tween-20 [TBST; 10 mm Tris HCl, 150 mm NaCl, and 0.05% Tween-20 (pH 8.0)] at room temperature for one hour, then probed overnight with the following primary antibodies (diluted in TBST): phospho-FAK (1:1000), total FAK (1:1000) from Cell Signaling (Danvers, MA); phospho-YES (1:1000); total-YES (1:500–1:1000); 4-HNE (1:700) from Abcam (Waltham, MA); and MDA clone -1F83 (1:200) from VWR (Radnor, PA). Nitrocellulose membranes were then incubated with horseradish peroxidase (HRP)-conjugated secondary antibodies from Cell Signaling (Danvers, MA) (1:2000) at room temperature for one hour, followed by incubation in enhanced chemiluminescence from Advansta (San Jose, CA) and imaged in the Amersham imager 600 (GE Healthcare). Membranes were reprobed for beta-actin (Cell Signaling, 1:1000) for ≥1 h at room temperature as a loading control.

### Cell Viability Assays

On Day 0, 300,000 – 400,000 cells of SUM44 and LCCTam were seeded in 6-well plates. Twenty-four hours later, on Day 1, cells were treated with control (DMSO), or Riluzole (10 μM), or a combination of Riluzole and Ferrostatin-1 (10 uM). On day 2, the cells were collected after a 24hr treatment period. The collected cells were stained with trypan blue and counted using the Countess II automated cell counter (Thermofisher). Data are analyzed using Prism 9 and presented as mean ± standard deviation (SD) of the ratio of the cell number of the treatment groups relative to the control for 3-4 independent biological assays.

### HCI-013 and HCI-013EI Patient Derived Xenograft (PDX) Experiments

All animal studies were ethically conducted in accordance with our approved Institutional Animal Care and Use Committee (IACUC) protocols #2018-0005 and #2018-0006. For the comparison of time to tumor formation between HCI-013 and HCI-013EI in the presence vs. absence of supplemental estrogen pellets, 5-6 week-old intact (non-ovariectomized) female non-obese diabetic, severe combined immunodeficient mice (NOD.CB17-Prkdcscid/NCrCrl, purchased from Charles River, Wilmington, MA) were orthotopically implanted into the right 4th mammary gland with a single 1-3 mm^3^ PDX fragment per mouse as follows: HCI-013EI (n=6); HCI-013EI+E2 (n=6); HCI-013 (n=6); HCI-013+E2 (n=6). “+E2” denotes co-implantation of a 1 mg estrogen pellet under the skin on the back between the shoulder blades. Mice were followed until measurable tumor development (by calipers), and data are presented as a survival plot with n=6 mice per group.

For the treatment study of HCI-013EI tumors, 5–6-week old intact (non-ovariectomized) female mice were orthotopically implanted into the right 4th mammary gland with a single 1-3 mm^3^ HCI-013EI PDX fragment per mouse without estrogen supplementation, then followed until tumors reached ~100 mm^3^ before enrollment to one of the four (4) treatment arms: Control (n=5); 25mg/kg Fulvestrant in castor oil SQ (once per week, n=5); 10 mg/kg Riluzole PO in corn oil (5 days per week, n=5); or the combination (n=5) for eight weeks. Mice were monitored for tumor growth (measured by calipers) and body weight twice per week. Tumor volumes were calculated by the modified ellipsoid formula V=1/2(XY^2^), where X is the longest axis, and Y is the longest perpendicular axis. Tumor volume data are presented as mean ± SEM for the number of mice per treatment group. The baseline measurement represents the measurement at the point of enrollment to a treatment group based on the *a priori* tumor volume as calculated above. The subsequent measurements are those taken while the mice are on treatment at the twice per week frequency. At the study’s conclusion, mice were humanely euthanized by approved AVMA guidelines. Tumors from the treatment study were resected, weighed, formalin-fixed, and paraffin-embedded.

### Standard Immunohistochemistry (IHC) Staining

Sections from formalin-fixed, paraffin-embedded tissues were deparaffinized with xylenes and rehydrated through a graded alcohol series. Heat-induced epitope retrieval (HIER) was performed by immersing the tissue sections at 98°C for 20 minutes in LowFlex (Dako #K8005). Staining was performed following the epitope retrieval process using VectaStain Kit from Vector Labs for cleaved Caspase-3 and horseradish peroxidase-labeled polymer from Dako (K4001) for PCNA. Slides were treated with 3% hydrogen peroxide and 10% normal goat serum for 10 minutes each and exposed to primary antibodies-1/120 for Caspase-3 and 1/1000 for PCNA Santa Cruz #sc56-for one hour at room temperature. Slides were then exposed to appropriate biotin-conjugated secondary antibodies, Vectastain ABC reagent, and DAB chromagen (Dako) for cleaved Caspase-3 and HRP labeled polymer and DAB chromagen (Dako) for PCNA. Slides were counterstained with Hematoxylin (Fisher, Harris Modified Hematoxylin), blued in 1% ammonium hydroxide, dehydrated, and mounted with Acrymount.

### IHC Imaging and Analysis

Slides were scanned at 40X magnification using the Aperio GT 450, an automated digital pathology slide scanner. The whole slide scans were viewed and analyzed with QuPath-0.3.0, open-source software used for bioimage analysis [37]. The images from the Caspase-3 and PCNA slides were separated into respective project groups, and a representative image from each group was analyzed. The corresponding analysis setting was then applied to the group to ensure uniformity across all the images. First, the default stain vector was selected to deconvolute the hematoxylin and DAB stains. Next, a region of interest (ROI) for analysis was selected; in this study, the entire tumor area was established as the region of interest for this analysis. Finally, the ROI was analyzed for positive stain detection, and the results (number positive per mm squared) were exported as a CSV file for statistical analysis using GraphPad Prism 9. Data for both PCNA and cleaved Caspase-3 are presented as the individual and median positive cells per mm^2^ for each treatment group.

### Multiplex Immunohistochemistry (mIHC) Staining

HCI-013+E2 (n=5) and HCI-013EI (n=4) tumors from mice in our PDX maintenance colony - independent from experimental animals described above - were resected, formalin-fixed, paraffin-embedded, then sectioned for staining on the Vectra3 multispectral imaging platform (Akoya Biosciences, Marlborough, MA) using OPAL chemistry. The slides were baked at 60°C, deparaffinized in xylene, rehydrated, washed in tap water, and incubated with 10% neutral buffered formalin for 20 minutes to increase tissue-slide retention. Epitope retrieval/microwave treatment (MWT) for all antibodies was performed by boiling slides in Antigen Retrieval buffer 6 (AR6 pH6; Akoya, AR6001KT). Protein blocking was performed using antibody diluent/blocking buffer (Akoya, ARD1001EA) for 10 minutes at room temperature. Primary antibody/OPAL dye pairings and incubation conditions for ER, PR, HER2, Ki67, and pan-cytokeratin staining are detailed in **Table 1**. MWT was performed to remove the primary and secondary antibodies between rounds of multiplex IHC. Multiplex IHC was finished with MWT, counterstained with spectral DAPI (Akoya FP1490) for 5 min, and mounted with ProLong Diamond Antifade (ThermoFisher, P36961). The order of antibody staining and the antibody/OPAL pairing was predetermined using general guidelines and the particular biology of the panel. General guidelines include spectrally separating co-localizing markers and separating spectrally adjacent dyes. Multiplex IHC was optimized by first performing singleplex IHC with the chosen antibody/OPAL dye pair to optimize signal intensity values and proper cellular expression, followed by optimizing the entire multiplex assay.

**Table 1:** Primary Antibody/OPAL Dye Pairings and Incubation Conditions.

|  | Antibody 1 | Antibody 2 | Antibody 3 | Antibody 4 | Antibody 5 |
| --- | --- | --- | --- | --- | --- |
| Antigen | PR | Ki67 | HER2 | ER alpha | panCK |
| Company | Agilent | Agilent | Agilent | Santa Cruz | Agilent |
| Cat# | M3569 | M7240 | A0485 | sc-8002 | M3515 |
| Species | Mouse | Mouse | Rabbit | Mouse | Mouse |
| Dilution | 1/50 | 1/50 | 1/200 | 1/50 | 1/300 |
| Incubation Time | 1 hr | overnight | 1 hr | overnight | 1 hr |
| Incubation Temp. | RT | 4°C | RT | 4°C | RT |
| Control Tissue | Breast Cancer | Tonsil | Breast Cancer | Breast Cancer | Breast Cancer |
| OPAL Fluor. | 650 | 520 | 620 | 570 | 690 |
| OPAL Conc. | 1/140 | 1/30 | 1/160 | 1/125 | 1/30 |
| Antigen Retrieval | AR6 | AR6 | AR6 | AR6 | AR6 |

### mIHC Imaging and Analysis

Slides were scanned at 10X magnification using the Vectra 3.0 Automated Quantitative Pathology Imaging System (PerkinElmer/Akoya). Whole slide scans were viewed with Phenochart (Perkin Elmer/Akoya), which allows for selecting high-powered images at 20X (resolution of 0.5m per pixel) for multispectral image capture. Multispectral images of each xenograft tissue specimen were captured in their entirety. Multispectral images were unmixed using spectral libraries built from images of single stained tissues for each reagent using the inForm Advanced Image Analysis software (inForm 2.4.6; PerkinElmer/Akoya). A selection of 10-15 representative multispectral images spanning all nine tissue sections was used to train the inForm software (tissue segmentation, cell segmentation, and phenotyping tools). All the settings applied to the training images were saved within an algorithm to allow the batch analysis of all the multispectral images particular to each panel. Data are presented as the overall mean ± SD of % marker positivity for all tumors.

### Fluorescence Lifetime Imaging (FLIM) Instrumentation

A modified Olympus FVMPERS (Waltham, MA) microscope equipped with a Spectra-Physics Insight X3 (Milpitas, CA) laser and FastFLIM (ISS, Champaign, IL) acquisition card were used to image the cancer samples. The samples were excited by two-photon excitation at 740 nm using a 20X air objective (LUCPLFLN 0.45NA, Olympus), and the emitted fluorescence was collected using the DIVER (Deep Imaging Via Enhanced Recovery) detector assembly equipped with a FastFLIM card for lifetime imaging. The pixel dwell time was fixed at 20 μs, and the field of view was 318.8 μm (Zoom =2X) at 256X256 pixels. 16 frames were integrated to increase signal-to-noise. The data from each pixel were recorded and analyzed using the SimFCS software (available from the Laboratory for Fluorescence Dynamics, University of California, Irvine, CA). The raster scanning was done using the Olympus software, and the images were collected using the FLIMBox/FastFLIM system in passive mode [38].

The samples (5 μm thick) were imaged using the homebuilt DIVER (Deep Imaging via Enhanced Recovery) microscope [39], a homebuilt modified detector based on an upright configuration. The details of this microscope have been described elsewhere [40,41]. Briefly, this microscope uses a forward detection scheme and a large area photon counting detector (R7600P-300, Hamamatsu), having a higher photon collection efficiency due to the large cone angle of detection. A combination of filters capable of separating the blue wavelength (400 – 500 nm) fluorescence was used for FLIM imaging of NADH [42]. The phasor plot is calibrated using Rhodamine 110 in water which has a mono-exponential lifetime of 4.0 ns.

### FLIM Phasor Analysis [38,43,44]

The fluorescence intensity decays collected at each pixel of the image were transformed to the Fourier space, and the phasor coordinates were calculated using the following relations:

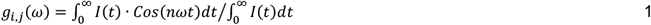

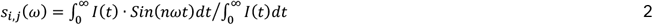

where *g_i,j_*(*ω*) and *s_i,j_*(*ω*) are the X and Y coordinates of the phasor plot, respectively, and *n* and *ω* are the harmonic numbers and the angular frequency of excitation, respectively. The transformed data were then plotted in the phasor plot so that the data from each pixel is transformed to a point in the phasor plot [45–47]. The fractional intensity distribution between free and protein-bound NADH was calculated based on a two-component analysis of the phasor plot [38,48] and then converted to concentration ratio based on the quantum yield of the two species [49]. The higher free/bound NADH ratio is representative of increased glycolysis [43].

### Primary Breast Tumor Explant Cultures

Patient-derived explants (PDEs) from five (5) ER+ primary breast tumors were processed and cultured as described in [50]. PDEs were treated with 100 nM Fulvestrant, 10 μM Riluzole, the combination, or solvent control (DMSO) for 48 hours before formalin fixation, paraffin embedding, sectioning, and staining for PCNA (1:1000, #sc-56, SCBT, Santa Cruz, CA), cleaved caspase 3 (1:300, #9661, Cell Signaling Technology, Danvers, MA), and Ki67 (1:500, #ab16667, Abcam, Cambridge, MA). Stained sections were then visualized and scored as described in [50]. Data are presented as change relative to vehicle (set to 0) for each PDE.

### Statistical Analysis

Statistical analyses were performed using GraphPad Prism 9.0 (San Diego, CA) at α≤0.05, except for Riluzole/Fulvestrant and Riluzole/4-hydroxytamoxifen combination experiments, which were analyzed by SynergyFinder [51]. Single-agent Riluzole experiments were analyzed by nonlinear regression ([inhibitor] vs. normalized response), and response to 10 μM Riluzole in endocrine therapy sensitive/resistant cell line pairs (SUM44 vs. LCCTam and MCF7 vs. LCC9) was compared by Mann-Whitney test.

Riluzole/Fulvestrant and Riluzole/4-hydroxytamoxifen combination experiments were analyzed using the Bliss, zero interaction potency (ZIP), and highest single agent (HSA) methods [52] in SynergyFinder. Cell cycle and Annexin V apoptosis assays were analyzed by two-way Analysis of Variance (ANOVA) followed by Sidak’s multiple comparisons tests. Cell viability assays for Riluzole Ferrostatin-1 were analyzed by two-way ANOVA followed by Tukey’s multiple comparisons test. Staining for each marker in primary breast tumor explant cultures was analyzed by one-sample t-test vs. 0 (vehicle). In the xenograft experiment comparing time to tumor formation between HCI-013 and HCI-013EI in the presence vs. absence of supplemental estrogen pellets, data were analyzed by log-rank (Mantel-Cox) test. In the xenograft experiment testing Fulvestrant, Riluzole, the combination, or control in HCI-013EI, tumor volume, and mouse body weight data were analyzed by mixed-effects analysis followed by Dunnett’s multiple comparisons tests at each time point vs. control. Tumor weight at the endpoint and PCNA and cleaved caspase-3 were analyzed by Browne-Forsyth and Welch ANOVA, followed by Dunnett’s T3 multiple comparison tests. Partial response (PR), stable disease (SD), and progressive disease (PD) were calculated using RECIST 1.1 criteria [53]. mIHC data were analyzed by the Mann-Whitney test.

## Results

Riluzole has shown anti-tumor activity in preclinical models of multiple cancers, including melanoma, glioblastoma, and breast cancer [25–31]. In several (but not all) of these reports, Riluzole-mediated growth inhibition is attributed to increased expression of metabotropic glutamate receptors (mGluRs, GRMs). We previously reported that acquisition of Tamoxifen resistance in a cellular model of invasive lobular breast cancer (ILC, [13]) is accompanied by the upregulation of GRM mRNA expression and growth inhibition by Riluzole [14]. Here, our goal was to test the efficacy of Riluzole more broadly, alone and in combination with multiple endocrine therapies, in a diverse set of ER+ *in vitro* and *in vivo* models enriched for ILC.

### Riluzole suppresses growth in ER+ breast cancer cell lines

We performed dose-response assays of Riluzole (33 nM to 100 μM) in four ILC- and two IDC-derived cell lines and the ER- non-transformed breast epithelial cell line MCF10A, using crystal violet staining as a proxy for total cell number [14] (**Figure 1A**). Nonlinear regression analysis calculated the Riluzole IC_50_ for all six cell lines to be ~10-100 μM, consistent with published studies in other malignancies. Direct comparison of growth inhibition by 10 μM Riluzole in three endocrine-responsive and -resistant cell line pairs (**Figure 1B**) confirmed [14] that the Tamoxifen-resistant ILC cell line LCCTam, and the MM134 LTED cells were significantly more responsive to Riluzole than their parental counterparts SUM44 and MM134 (Mann-Whitney test, **p= 0.002 & 0.0043 respectively). This was not the case for the MCF7/LCC9 IDC cell line pair [54], in which MCF7 cells showed greater Riluzole-mediated growth inhibition (*p=0.024) than Fulvestrant-resistant/Tamoxifen-cross-resistant LCC9 cells. However, MCF10A non-transformed cells were not growth inhibited by 10 μM Riluzole vs. DMSO control.

**Figure 1.**
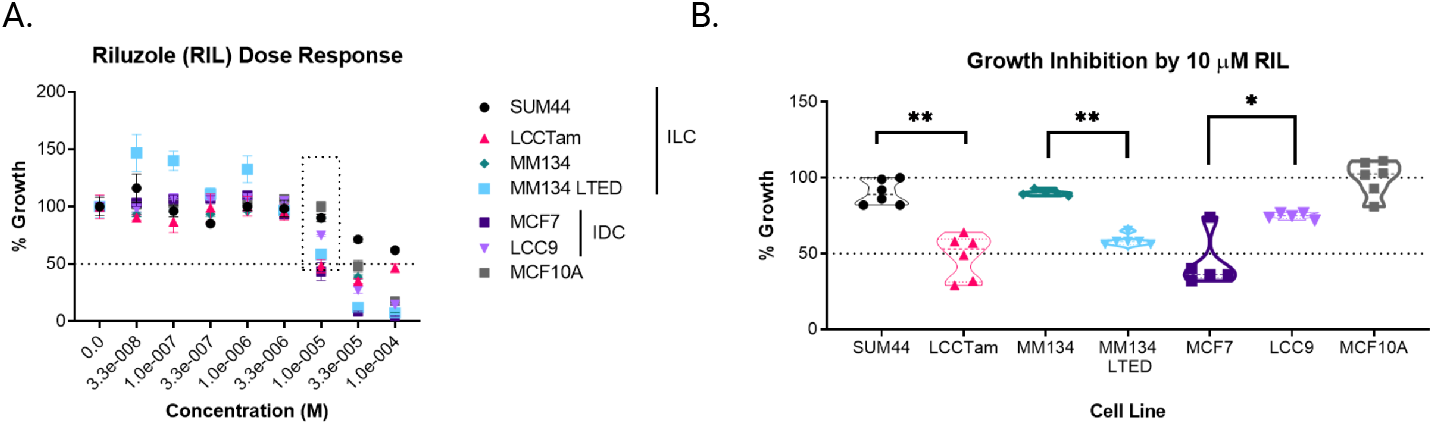
Growth suppression of ER+ breast cancer cell lines by Riluzole. **A**, Cells seeded in 96-well plates were treated with the indicated concentrations of Riluzole (RIL, 33 nM to 100 μM) or DMSO control for 7-8 days prior to staining with crystal violet. Data are presented as mean % growth ± standard error of the mean (SEM) of % growth (vehicle = 100%) for 5-6 technical replicates and represent 2-4 independent biological assays. The dotted line box indicates data re-graphed in panel B. Data were analyzed by nonlinear regression ([inhibitor] vs. normalized response), yielding the following IC_50_ [M] estimates: SUM44, 1.27e-4; LCCTam, 2.13e-5; MM134, 2.73e-5; MM134 LTED, 1.209e-5; MCF7, 1.09e-5; LCC9, 2.23e-5; MCF10A, 4.33e-5. **B**, Relative response to 10 μM RIL re-graphed from panel A (dotted line box). Data are presented as median % growth with upper/lower quartiles of % growth (vehicle = 100%) for 5-6 technical replicates and represent 2-4 independent biological assays. For the SUM44/LCCTam, MM134/MM134 LTED, and MCF7/LCC9 cell line pairs, data were compared by the Mann-Whitney test. **p=0.002, **p = 0.0043, and *p=0.024 respectively. Dashed lines denote 50% (panels A and B) and 100% growth inhibition (panel B).

The presence vs. absence of steroid hormones and estrogenic compounds in growth media (e.g., phenol red, serum) can influence the response of ER+ cell lines to growth inhibition by small molecules. The SUM44/LCCTam cell line pair is cultured in serum-free media, but a phenol red-containing base (IMEM, 10 mg/L), while MCF7 and MM134 cells are cultured in phenol red-containing, IMEM supplemented with 5%FBS. LCC9 cells and MM134 LTED are maintained in hormone-replete conditions. As such, experiments presented in Figure 1 performed under hormone-replete conditions were repeated under hormone-deprived conditions (**Figure S1** [55]). While individual differences within cell lines were observed, hormone deprivation - reduced phenol red media for SUM44/LCCTam (Ham’s F12, 1.2 mg/L) or phenol red-free IMEM supplemented with 5% CCS for MCF7 and MM134 - did not consistently enhance or impair Riluzole-mediated growth inhibition.

### Riluzole induces a histologic subtype-associated cell cycle arrest

To corroborate the cell proliferation assay results, we tested Riluzole’s effect on cell cycle progression (**Figure 2**). All ILC cell lines (including BCK4, a third model of ER+ ILC [32], **Figure S2** [55]) showed a significant accumulation of cells in the G2-M phase (two-way ANOVA followed by Sidak’s multiple comparisons test, see figure legends). However, both IDC-derived cell lines showed a significant accumulation of cells in the G0-G1 phase, while non-transformed MCF10A cells showed no significant cell cycle arrest in response to Riluzole. Together with the results presented in Figures 1 and S1, these data suggest that while all ER+ breast cell lines tested are growth inhibited by Riluzole, ILC cells preferentially undergo G2-M arrest while IDC cells arrest in G0-G1.

**Figure 2.**
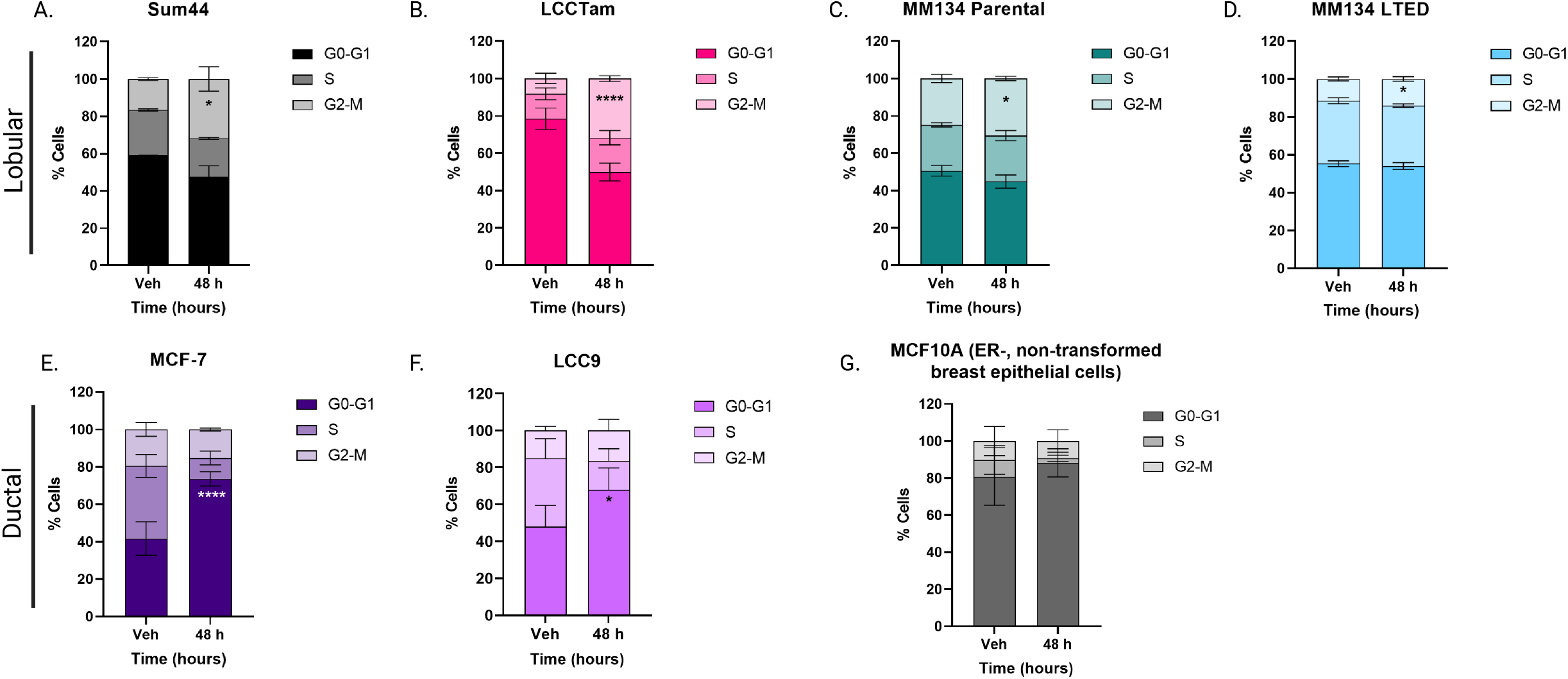
Riluzole induces a histologic subtype-associated cell cycle arrest. Cells seeded in 6-well plates were treated with 10 μM Riluzole or DMSO control (vehicle, Veh) for the indicated times prior to collection, fixation, staining, and cell cycle analysis. Data are presented as mean % cells ± SD for 3-4 independent biological assays and analyzed by two-way ANOVA followed by either Sidak’s (single time point) or Dunnett’s (multiple time points) multiple comparisons tests. SUM44: *p = 0.018. LCCTam: ****p < 0.0001. MM134: *p = 0.011. MM134 LTED: *p = 0.05. MCF7: ****p < 0.0001. LCC9: *p = 0.015. MCF10A: not significant.

### Riluzole inhibits phosphorylation of pro-survival signaling molecules and induces apoptosis and ferroptosis

To identify molecular signaling events accompanying Riluzole-mediated growth inhibition in the SUM44/LCCTam cell line pair, we used the Human Phospho-Kinase Proteome Profiler™ Array to detect changes in 43 phosphorylation sites across 40 different kinases or substrates (**Figure 3A**). In SUM44 cells, Riluzole reduced phosphorylation of mutant p53 [56] (S92 and S392), and Akt T308. In LCCTam cells, Riluzole reduced phosphorylation of markedly more sites in kinases and substrates with several significantly enriched ontology clusters, including Signaling by Receptor Tyrosine Kinases and Cytokine Signaling in Immune System (**Figure S3A** [55]). In addition, notable inhibition of Akt/mTOR (Akt S437, TOR S2448, PRAS40 T246), CREB (Msk S376/360, CREB S133) and Src/Fak (Lyn Y397, Yes Y426, Fak Y397) signaling pathways were observed specifically in LCCTam cells. Prior studies in melanoma and glioblastoma have shown that Riluzole inhibits Akt phosphorylation and that Riluzole combined with mTOR inhibition can synergistically decrease xenograft growth [26,57]. However, to our knowledge, inhibition of Src/Fak family kinases by Riluzole has not been previously reported. Therefore, to further validate the inhibition of selected Src/Fak kinases observed from the Human Phospho-Kinase Proteome Profiler™ Array, we performed a western blot analysis on SUM44/LCCTam cell line treated with Riluzole for several time points (6,12, 24, and 48 hours). The results show markedly higher baseline Fak Y397 phosphorylation and confirmed reduced expression and phosphorylation of Fak Y397 in the LCCTam cells versus no change to a slight increase in Fak phosphorylation in SUM44 cells (**Figure 3B, 3C**). However, we could not validate a change in Yes expression or phosphorylation at Y426 (**Figure S3B** [55]).

**Figure 3.**
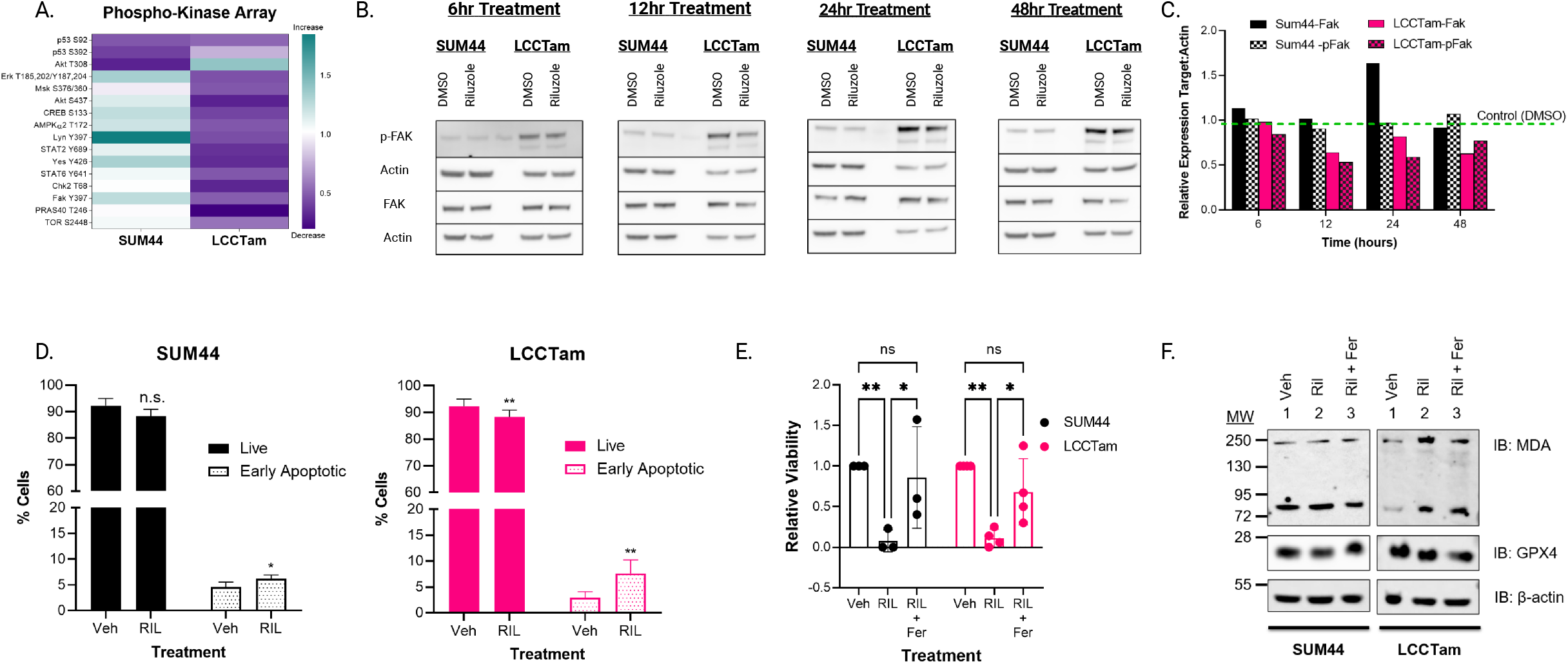
Riluzole inhibits phosphorylation of pro-survival signaling molecules and induces apoptosis and ferroptosis. **A**, Cells seeded in 6-well plates were treated with 10 μM Riluzole or DMSO control (vehicle, Veh) for 2 days prior to collection, lysis, and processing, then assayed using the Human Phospho-Kinase Proteome Profiler™ Array. A ratio of background-corrected intensity values for targets (phospho-kinase spots) to references (control spots) was created for each condition (DMSO and Riluzole) within each cell line. Data are presented as the geometric mean of the Riluzole: DMSO ratio for 2 technical replicates from a single experiment. **B**, Sum44, and LCCTam cells seeded in 6-well plates were treated with 10 μM Riluzole or DMSO control for several time points (6,12, 24, and 48 hours). After which, cells were collected, lysed, and western blot analysis was performed to test for expression and phosphorylation of Fak Y397 phosphorylation. The data are presented as images showing expression levels. **C**, Quantification analysis of Fak and p-Fak protein band density from western blot in Figure 3B. **D**, Cells seeded in 6-well plates were treated with 10 μM Riluzole or DMSO control (vehicle, Veh) for 2 days prior to staining with Annexin V and PI. The percent of live cells (PI^-^, annexin V^-^) and early apoptotic (PI^-^, annexin V^+^) cells are shown. Data are presented as mean % cells ± SD for 3 (SUM44) or 4 (LCCTam) independent biological assays and analyzed by two-way ANOVA followed by Sidak’s multiple comparisons test (*p =0.021, **p = 0.04 (live) and **p= 0.011). **E**, Cells of SUM44 and LCCTam were seeded in 6-well plates. Twenty-four hours later, cells were treated with control (DMSO), Riluzole (10 μM), or a combination of Riluzole and Ferrostatin-1 (10 uM). 24h after treatment, the cells were collected, stained with trypan blue, and counted. Data are presented as mean ± standard deviation (SD) of the ratio of the cell number of the treatment groups relative to the control for 3-4 independent biological assays and analyzed by two-way ANOVA followed by Tukey’s multiple comparison test (Sum44 – (**p = 0.005, *p = 0.016), LCCTam – (**p = 0.002, *p = 0.043)). **F**, Sum44, and LCCTam cells seeded in 6-well plates were treated with control (DMSO), Riluzole (10 μM), or a combination of Riluzole and Ferrostatin-1(10 μM) for 48hr. Post-treatment, cells were collected, lysed, and western blot analysis was performed to test for expression of Malondialdehyde (MDA). The data are presented as images showing expression levels.

We then performed Annexin V assays to measure the effect of Riluzole on apoptosis in the SUM44/LCCTam cell line pair. LCCTam cells showed a significant increase in the percent of cells in early apoptosis when treated with Riluzole (**Figure 3D** and **S3C** [55], two-way ANOVA followed by Sidak’s multiple comparisons tests, **p<0.001). However, there was a less robust increase in early apoptotic SUM44 cells (two-way ANOVA followed by Sidak’s multiple comparisons test, *p<0.0211). These data are consistent with those presented in Figure 1B, where LCCTam cells were significantly more growth-inhibited by Riluzole than SUM44 cells.

In addition to apoptosis, the iron-dependent cell death mechanism of ferroptosis could be relevant to Riluzole action in these cells. For example, inhibition of the PI3K-Akt-mTOR pathway has been previously implicated in this form of cell death [58]. Furthermore, Fak signaling downstream of the glutamate/cystine antiporter SLC7A11 or X_c_^-^, which can be inhibited by Riluzole, has also been implicated in ferroptosis [59]. To explore the possibility of ferroptotic cell death, we performed a viability assay after treating Sum44 and LCCTam cells with Vehicle (DMSO), or Riluzole or a combination of Riluzole and Ferrostatin-1 (inhibitor of Ferroptosis). The results showed that Riluzole reduces cell viability in both SUM44 and LCCTam, and the addition of Ferrostatin-1 reverses the observed reduction (**Figure 3E**). To substantiate this observation, we performed a western blot analysis on the SUM44/LCCTam cell line pair treated with vehicle (DMSO) or Riluzole, or a combination of Riluzole and Ferrostatin for 48hrs, then probed for Malondialdehyde (MDA) and 4-Hydroxynonenal (4-HNE) – which are both by-products of ferroptosis. MDA levels were increased in LCCTam cells treated for 48 hr with Riluzole. Conversely, ferrosatin-1 decreased the Riluzole-induced MDA (**Figure 3F**). On the other hand, Riluzole slightly increased the levels of 4-HNE in LCCTam cells, whereas the combined treatment of Riluzole and Ferrostatin-1 reduced the levels of Riluzole-induced 4-HNE (**Figure S3D** [55]). Altogether, the Riluzole induction of lipid peroxidation products suggests Riluzole-induced ferroptosis in LCCTam.

### Riluzole, in combination with endocrine therapies, leads to additive suppression of ER+ breast cancer cell line growth

Endocrine therapies ranging from SERMs and SERDs to aromatase inhibitors represent the standard of care for the clinical management of ER+ breast cancers [2]. Therefore, we tested Riluzole’s activity in combination with the SERD Fulvestrant or SERM Tamoxifen (4-hydroxytamoxifen) in ILC- and IDC-derived ER+ breast cancer cell lines and the ER- non-transformed breast epithelial cell line MCF10A as a negative control. These experiments were conducted under hormone-replete conditions. To determine the possible relational effect of the drug combinations, we used SynergyFinder, a web-based tool for interactive analysis and visualization of multi-drug and multi-dose response data [51]. Based on the synergy finder scoring, the combination of Fulvestrant and Riluzole showed additive benefits in nearly all tested cell lines (**Figure 4** and **S4A** [55]). The representative synergy map of the bliss model highlights the synergistic and antagonistic dose regions in red and green, respectively, and the overall synergy score indicated at the top (**Figure 4A**, >10 = synergy, 10 to - 10 = additive, <-10 = antagonistic). Examination of the other synergy models provided similar synergy scores to the bliss model in Figure 4A, which supports the notion that the combination of Fulvestrant and Riluzole is additive (**Figure 4B**). The synergy analysis of the combination of Riluzole and Tamoxifen resulted in scores that indicated an additive interaction in all the cell lines except for MM134 LTED and MCF10A (**Figure 4B**). MM134 LTED cells were only inhibited by the lowest concentration of 4HT and not by the higher concentrations (**Figure S4B** [55]). On the other hand, Riluzole significantly inhibited growth, therefore, the drugs’ opposing effects likely account for the combination’s antagonistic effect. Despite these exceptions, these data suggest that the combination of endocrine therapy and Riluzole, in most cases, can additively suppress the growth of a variety of ER+ breast cancer cell line models.

**Figure 4.**
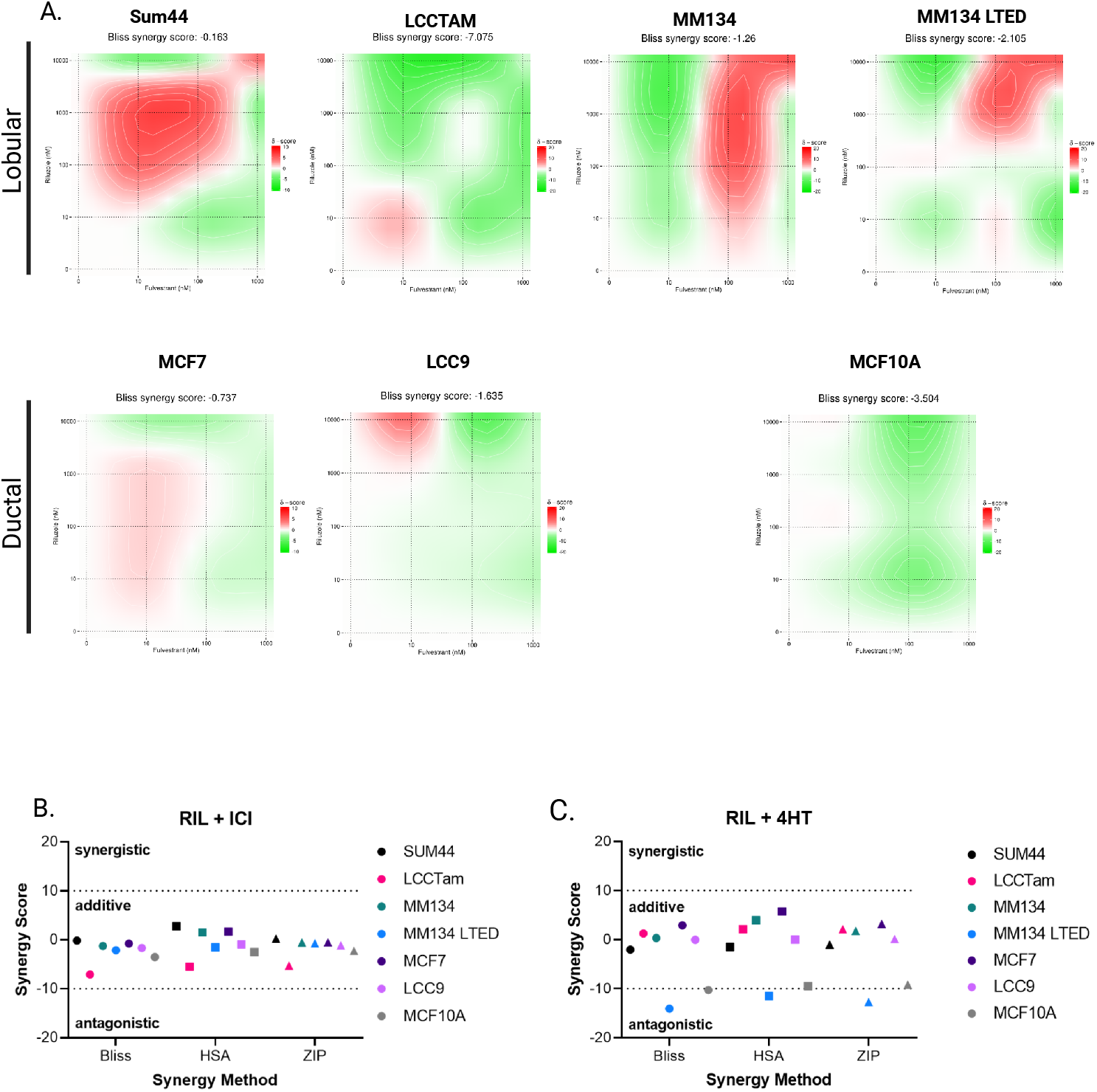
Additive suppression of ER+ breast cancer cell line growth by Riluzole in combination with endocrine therapies. **A**, Cells seeded in 96-well plates were treated with 1 μM Fulvestrant, 10 μM Riluzole (RIL), the combination, or DMSO control (vehicle, Veh) for 7-8 days prior to staining with crystal violet. Data are processed as the mean % growth for 5-6 technical replicates and represent 2-4 independent biological assays. The mean % growth data of a representative single technical replicate was then used to create a combination matrix used in SynergyFinder, and the results were presented as 2D surface plots. The SynergyFinder scores are shown at the top of the plots, highlighting the level of synergy. **B** and **C**, Graphical representation of the synergy scores from the Riluzole – Fulvestrant (**B**) and Riluzole – 4HT (**C**) combination using 3 models – Bliss, HSA, and ZIP.

### Single-agent Riluzole inhibits tumor growth *in vivo*, but the combination with Fulvestrant is not better than Fulvestrant alone in the HCI-013EI ILC PDX model

PDXs are an important, clinically relevant alternative to 2D culture models for pre-clinical testing of combination therapies. The HCI-013 PDX model was established from a 53-year-old woman with metastatic, multi-therapy-resistant ER+/PR+/HER2-ILC by serial passage through intact female NOD scid gamma (NSG) mice supplemented with a 1 mg estrogen (E2, 17β-estradiol) pellet [15]. The HCI-013EI (estrogen-independent) variant was established by two weeks of *in vitro* culture of cells from HCI-013 tumors under hormone-deprived conditions, then reimplanted into intact female NSG mice without estrogen supplementation [60]. Both models harbor the clinically relevant *ESR1* activating mutation Y537S, with the HCI-013EI variant reported as having a more abundant variant allele fraction of Y537S.

To directly compare the responsiveness to, and dependency on, supplemental estrogen of HCI-013 vs. HCI-013EI, six (6) 5-6 week-old intact severe combined immunodeficient (SCID) female mice per group were orthotopically implanted with a single 1-3 mm^3^ PDX fragment, then tumor growth and development was monitored (**Figure 5A**). In the presence of supplemental estrogen pellets, HCI-013 and HCI-013EI exhibited a 100% tumor take rate, with a median time to tumor formation of 23.5 and 26.5 days, respectively. However, in the absence of supplemental estrogen pellets, HCI-013 PDX fragments were unable to form tumors out to 113 days post-implantation, and HCI-013EI PDX fragments exhibited a 50% tumor take rate, with a median time to tumor formation of 39 days (log-rank Mantel-Cox test, ***p=0.0007). These data suggest that supplemental estrogen is necessary for HCI-013 tumor formation and beneficial but not necessary for HCI-013EI tumor formation in SCID mice.

**Figure 5.**
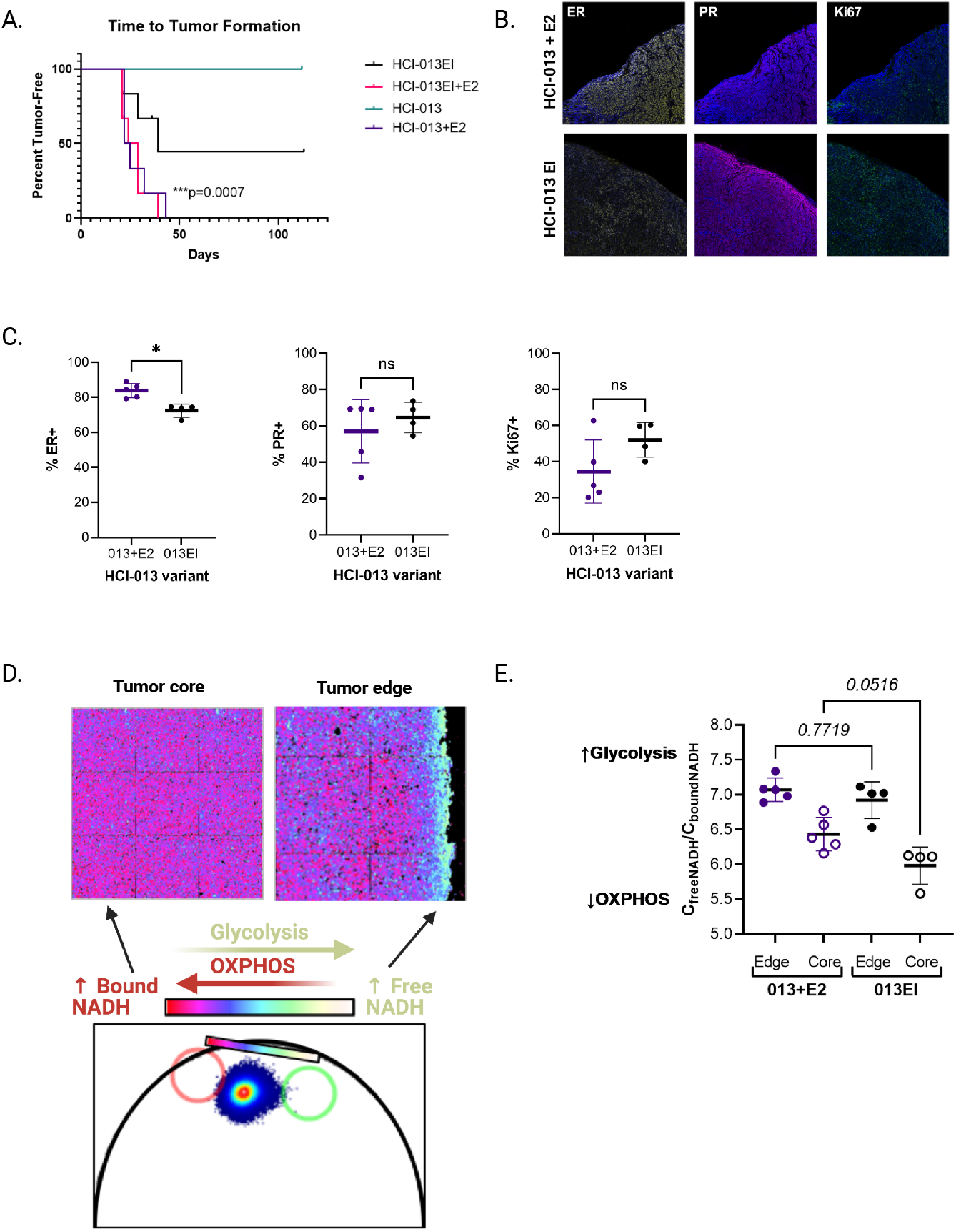
Characterization of tumor growth, hormone receptor expression, and metabolic state for HCI-013 vs. HCI-013EI ILC PDX models. **A**, Tumor latency for HCI-013 and HCI-013EI patient-derived xenografts (PDXs) with or without estrogen (E2) supplementation. Six (6) mice per group were orthotopically implanted with a 1-3 mm^3^ PDX fragment, then followed until measurable tumor development (by calipers). Data are presented as percent tumor free and were analyzed by log-rank (Mantel-Cox) test. ***p=0.0007. **B**, Representative images of ER, PR, and Ki67 staining from HCI-013+E2 and HCI-013EI tumors. **C**, Semi-quantitative analysis of multiplex IHC (mIHC) staining for ER, PR, and Ki67 from HCI-013+E2 (n=5) and HCI-013EI (n=4) tumors from mice independent of those for whom tumor latency is shown in panel A. Data are presented as overall mean ± SD of % marker positivity for 5-13 fields per tumor. **D**, Representative schematic of fluorescence lifetime imaging microscopy (FLIM) analysis of cellular metabolism at the tumor core and edge. Images are pseudo-colored based on the phasor plot (below) where more protein bound and more free NADH phasor positions are indicated by red and cyan circles, respectively, and the color scheme chosen reflects more bound NADH in purple and more free NADH in cyan. **E**, Quantification of FLIM in HCI-013+E2 vs. HCI-013EI tumor cores and edges. Tumor edges were strongly glycolytic in both HCI-013+E2 and HCI-013EI, but tumor cores were preferentially in an oxidative phosphorylated state in HCI-013EI.The data were analyzed by one-way ANOVA (p < 0.0001) followed by Tukey’s multiple comparisons test. Each symbol indicates the mean C_free NADH_/C_bound NADH_ for 16 fields of view of the tumor edge or core in an individual tumor (n=5 tumors per PDX line).

We characterized an independent set of HCI-013+E2 (estrogen supplemented, n=5) and HCI-013EI (not estrogen supplemented, n=4) tumors with respect to hormone receptor (ER and PR), HER2, and proliferative marker Ki67 expression using Opal chemistries on the Vectra3 multispectral imaging platform (**Figures 5B and 5C**). This approach captured heterogeneity in marker expression between and within tumors. Overall, percent ER positivity (% ER+) was significantly lower in HCI-013EI vs. HCI-013+E2 tumors (Mann-Whitney test, *p=0.032), consistent with a prior report that Y537S mutant ER protein expression can be lower than wild type ER [61]. However, overall percent PR and Ki67 positivity were not significantly different between these PDX variants. No HER2 staining was detected (data not shown).

An important feature of endocrine-resistant breast cancer is dysregulated metabolism, with published studies showing increased dependency on glutamine [62] and other amino acids [63]. Advanced imaging techniques like fluorescence lifetime imaging (FLIM) take advantage of the natural autofluorescence of biomolecules, including the reduced form of nicotinamide adenine dinucleotide (NADH, a key output of cellular metabolism). FLIM coupled with phasor analysis permits resolution of bound vs. free NADH, which correlates with oxidative phosphorylation vs. glycolytic metabolism, respectively [38,64]. Using FLIM, we examined the cellular metabolism of HCI-013+E2 and HCI-013EI tumors. We observed that cells were mainly glycolytic at the edge of both HCI-013+E2 and HCI-013EI tumors. However, cells at the core of HCI-013EI tumors were preferentially in an oxidative phosphorylated state as opposed to a more glycolytic state observed in cells within the core of the HCI-013+E2 tumors (**Figures 5D and 5E**, one-way ANOVA (p < 0.0001) followed by Tukey’s multiple comparisons test (p=0.0516).

We selected the HCI-013EI (not estrogen-supplemented) PDX variant to test the anti-tumor activity of Fulvestrant, Riluzole, or the combination relative to vehicle control. Forty-eight (48) 5-6 week-old intact SCID female mice were orthotopically implanted with a single 1-3 mm^3^ HCI-013EI PDX fragment without E2 supplementation, then followed until tumors reached ~100 mm^3^ before enrollment to one of four (4) treatment arms: control (n=5), Fulvestrant (n=5), Riluzole (n=5), or the combination (n=5) (**Figures 6A and S5A** [55]). Relative to control, single-agent Riluzole, Fulvestrant, and the combination significantly slowed the tumor volume. However, the level of tumor growth inhibition relative to control varied among the three groups. The Riluzole group showed about 50% inhibition, whereas similar effects but greater inhibition (~90%) were observed in the Fulvestrant and combination groups. (**Figure 6A**, mixed effect analysis followed by Tukey’s multiple comparisons tests). Analysis of tumor weight at the endpoint for the treatment groups further supported the observed difference in tumor volume. The mean weights of the Fulvestrant, Riluzole, and Combination groups were lower than the control group. However, only the Fulvestrant and combination groups showed statistically significant differences **(Figure 6B**, Browne-Forsyth and Welch ANOVA followed by Dunnett’s T3 multiple comparisons tests) and were not different from each other. Analysis of relative tumor size at endpoint according to RECIST 1.1 criteria [53] shows that 2 of 5 tumors in the Fulvestrant group and 3 of 5 in the combination group achieved partial response (PR) (**Figure 6C**). We then performed immunohistochemistry to stain for proliferating cell nuclear antigen (PCNA) and Caspase-3 as a proxy for proliferation and apoptosis, respectively. Although not significant, the mean positive cells per mm^3^ of Caspase-3 for the Fulvestrant, Riluzole, and combination group were each higher than the control group (**Figure 6D**). For the PCNA staining, expectedly, the Fulvestrant and combination group had lower mean positive cells per mm^3^. However, surprisingly, the mean positive stained cells per mm^3^ for the Riluzole group was higher than the control group (**Figure 6E**). Finally, analysis of mouse body weights between the treatment groups showed no significant differences. As seen in **Figure S5B** [55] the slope of the graphs for each treatment group is close to zero. Altogether, these data show that single-agent Riluzole has a significant inhibitory effect on HCI-013EI tumor volume, and with Fulvestrant already highly effective against this PDX model, combination treatment does not provide additional benefit.

**Figure 6.**
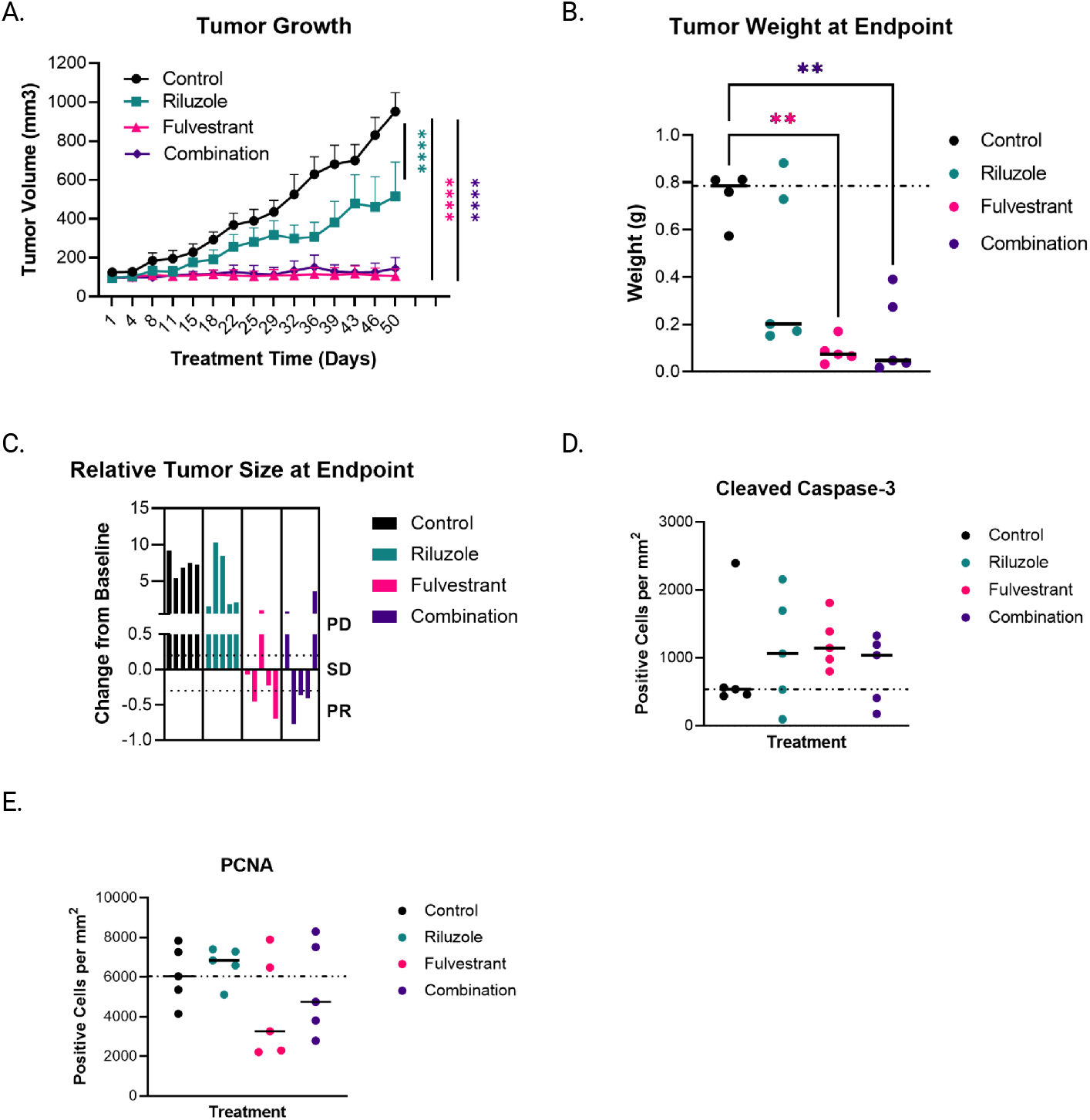
Single-agent Riluzole inhibits tumor growth in vivo, but the combination with Fulvestrant is not better than Fulvestrant alone in the HCI-013EI ILC PDX model. **A**, Forty-eight (48) mice were orthotopically implanted with a 1-3 mm^3^ HCI-013EI PDX fragment without E2 supplementation, then followed until tumors reached ~100 mm^3^ before enrollment to one of four (4) treatment arms: control (n=5), Fulvestrant (n=5), Riluzole (n=5), or the combination (n=5). Mice were monitored for tumor growth (measured by calipers) and body weight twice per week. Data are presented as mean tumor volume ± SEM, and were analyzed by mixed-effects analysis followed by Dunnett’s multiple comparisons tests at each timepoint vs. control. **B**, At the end of the study, tumors were collected and weighed. The graph illustrates the summary of the collected data, which were analyzed using Browne-Forsyth and Welch ANOVA followed by Dunnett’s T3 multiple comparisons tests. **C**, Graph showing relative tumor size at endpoint according to RECIST 1.1 criteria. It shows that 2 of 5 tumors in the Fulvestrant group and 3 of 5 in the combination group achieved partial response (PR). **D** and **E**, The tumors collected from each treatment group were formalin-fixed, paraffin-embedded, sectioned, and stained with proliferating cell nuclear antigen (PCNA) and Caspase-3 by IHC. These antibodies served as a proxy for proliferation and apoptosis, respectively. The stained samples were analyzed, and the data were presented graphically in **D** for PCNA, and **E** for Caspase-3.

### Riluzole plus Fulvestrant significantly inhibits proliferation in primary breast tumor explant cultures

Patient-derived explants (PDEs) provide another pre-clinical strategy to test combination therapies. These short-term cultures of surgical samples maintain the local tumor microenvironment and capture inter-person heterogeneity [50]. We tested the efficacy of Fulvestrant, Riluzole, or the combination vs. solvent control (DMSO) in five (5) PDEs from ER+/PR+/HER2-negative primary tumors (**Figure 7A**). Using PCNA staining as a proxy for cell proliferation, the combination of Riluzole plus Fulvestrant significantly reduced PCNA (**Figures 7B and 7C**, one-sample t-test vs. 0 (Vehicle), *p=0.013 Vehicle vs. Combination), with 4 of 5 PDEs showing better growth inhibition by the combination than either drug alone and the greatest effect seen in the ILC PDE. Staining for cleaved caspase 3 suggested a modest induction of apoptosis by either drug alone or the combination in some of the PDEs (**Figure S4A** [55]), but this was not statistically significant. Together with the results presented in Figures 5, 6, and accompanying supplementary figures, these data suggest that combining Fulvestrant and Riluzole may offer improved therapeutic benefits in some ER+ breast cancers.

**Figure 7.**
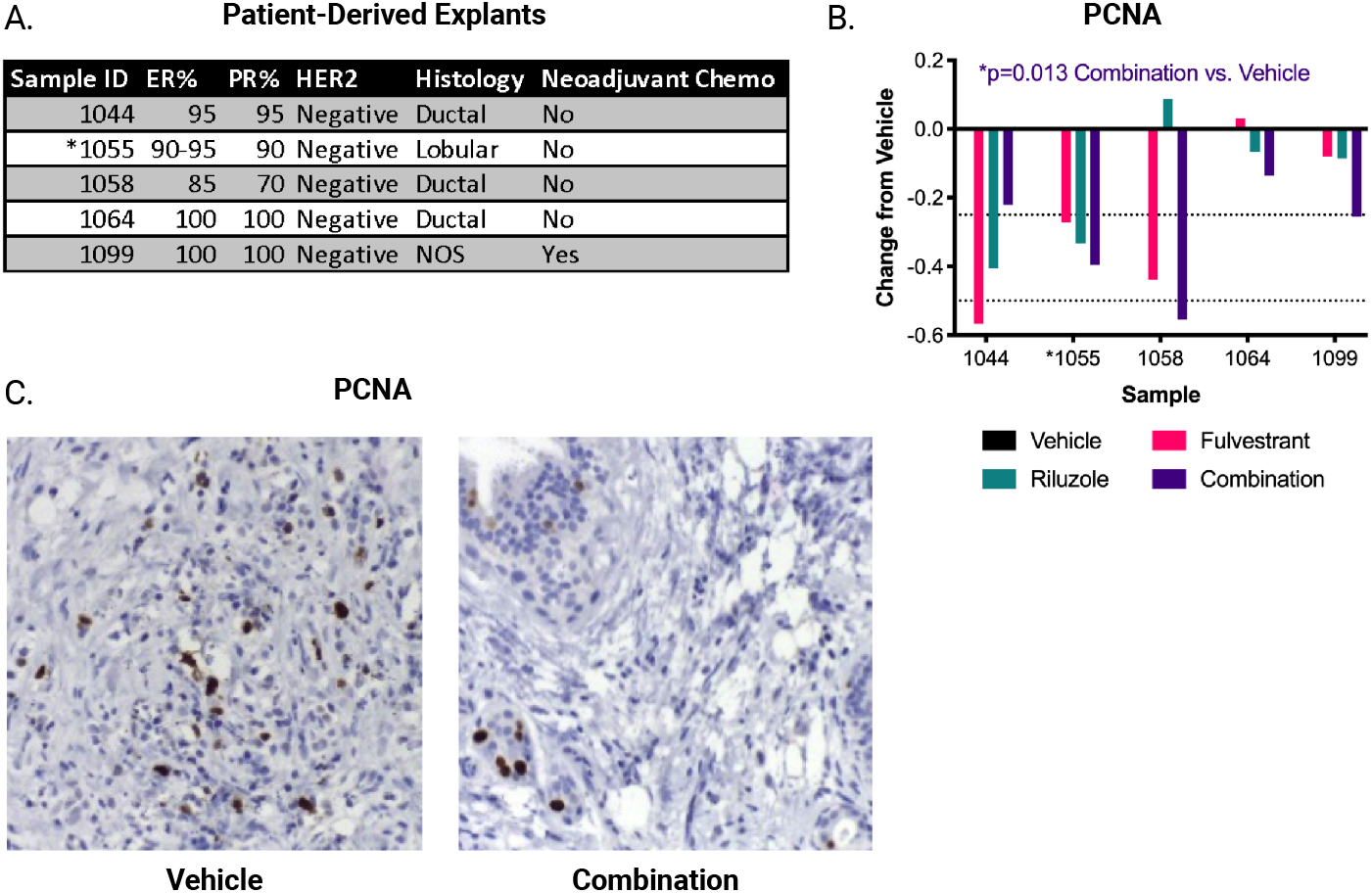
Riluzole plus Fulvestrant significantly inhibits proliferation in primary breast tumor explant cultures. **A**, Pathologic data for five (5) patient-derived explants (PDEs). ER, PR, and Ki67% are from the initial surgical specimen, and NOS = not otherwise specified. *denotes the PDE for which representative images are shown in panel C. **B**, PDEs were treated with 100 nM Fulvestrant, 10 μM Riluzole, the combination, or DMSO control (vehicle) for 2 days prior to formalin fixation, paraffin embedding, sectioning, and staining for PCNA by IHC. Data are presented as change relative to vehicle (set to 0) for each explant and analyzed by one-sample t-test vs. 0 (vehicle). *p=0.013 Vehicle vs. Combination. *denotes the PDE for which representative images are shown in panel C. **C**, Representative images of PCNA and Caspase-3 staining from PDE #1055 (ILC).

## Discussion

We tested the efficacy of Riluzole, alone and in combination with multiple endocrine therapies, in a diverse set of ER+ *in vitro* and *in vivo* models enriched for ILC. Single-agent Riluzole suppressed the growth of ER+ ILC and IDC cell lines *in vitro*, by inducing a histologic subtype-associated cell cycle arrest (G0-G1 for IDC, G2-M for ILC). In the Tamoxifen-resistant ILC-derived LCCTam model, Riluzole induced apoptosis and reduced phosphorylation of multiple pro-survival signaling molecules, including Akt/mTOR, CREB, and Src/Fak family kinases. Furthermore, in combination with either Fulvestrant or 4-hydroxytamoxifen, Riluzole additively suppressed ER+ breast cancer cell growth *in vitro*. In addition, single-agent Riluzole inhibited HCI-013EI ILC PDX growth *in vivo*, and the combination of Riluzole plus Fulvestrant significantly reduced proliferation in primary breast tumor explant cultures.

The increased combinatorial efficacy of Riluzole and Fulvestrant we observe across diverse cell line models of ER+ breast cancer *in vitro* is recapitulated *ex vivo* using PDEs. In the HCI-013EI PDX experiment, tumor volume after ~7 weeks of treatment was not significantly different in combination-vs. Fulvestrant-treatment groups, although single-agent Riluzole had significant activity. We attribute this partly to the very strong response of this PDX to single-agent Fulvestrant as seen in this experiment (~90% inhibition) and reported by others [45,46]. In the PDE experiment, the combination of Riluzole and Fulvestrant was highly effective, with 80% of primary tumor explants (4/5) showing significant growth inhibition by the combination as measured by a reduction in PCNA (Figure 7). Improved combinatorial efficacy in the PDEs may also be due to the lower concentration of Fulvestrant used in these studies (100 nM). Additionally, Riluzole bioavailability is variable, leading to mixed efficacy in preclinical and clinical studies. For example, in preclinical studies of triple-negative breast cancer [30] and glioblastoma [26], single-agent Riluzole does not have significant anti-tumor activity *in vivo*, whereas in our study and preclinical studies of melanoma [24,25,65] it does. However, in Figure 6B-C, two of five Riluzole-treated tumors showed no reduction in tumor weight or relative tumor size versus control-treated tumors, which could be due to inconsistent bioavailability. Serum levels of the drug vary widely in ALS patients receiving the drug, and in a phase II trial for advanced melanoma, circulating Riluzole concentrations had marked inter-patient variability [66]. The Riluzole pro-drug troriluzole [67] appears to offer better bioavailability by reducing first-pass metabolism by CYP1A2, leading to sustained ~0.3-1.8 μM plasma concentrations of active drug in a recent phase Ib study that combined troriluzole (BHV-4157) with nivolumab in advanced solid tumors [68]. These and other Riluzole analogs will be important to explore alone and in combination with endocrine therapy in ER+ breast cancer.

In the Tamoxifen-resistant ILC-derived LCCTam model, Riluzole induces apoptosis and ferroptosis concomitant with reduced phosphorylation of multiple pro-survival signaling molecules. That these include components of the Akt/mTOR signaling pathway (Akt S437, TOR S2448, PRAS40 T246) is not surprising since Riluzole has been shown to inhibit Akt phosphorylation and synergize with mTOR inhibition in melanoma and glioblastoma models [26,57]. However, Src/Fak kinase family members (e.g., Yes and Fak) have not, to our knowledge, been previously implicated in Riluzole action. In vehicle-treated LCCTam cells, baseline phosphorylation of Fak Y397 is markedly increased compared to vehicle-treated SUM44 cells (Figure 3B). At the same time, Riluzole treatment reduced total and phosphorylated protein levels of Fak in only the LCCTam cells (Figure 3B-C). Multiple Src/Fak kinase family members play critical roles in cell survival, invasion, and migration. Additionally, Fak has been previously implicated in endocrine therapy resistance [69,70]. Being functionally E-cadherin-negative, ILC is highly resistant to anoikis (a form of cell death induced by extracellular matrix detachment) [71] and dependent upon a rewired actin cytoskeleton and constitutive actomyosin contractility (reviewed in [72]). Consistent with this, expression of activated Src and Fak – both of which can drive resistance to anoikis [73] – are significantly higher in ILC vs. atypical lobular hyperplasia [74]. In parallel, ILC may exhibit increased sensitivity to ferroptosis inducers. Ferroptosis is cell density-dependent, with cells cultured in low-density conditions with fewer cell-cell contacts (a defining feature of ILC) being highly vulnerable to ferroptosis caused by inhibition of glutathione peroxidase 4 (GPX4, [75]). On the other hand, E-cadherin-mediated intercellular interactions can suppress ferroptosis via activation of neurofibromin 2 (NF2, [76]). The specific contribution of Fak to Riluzole-mediated growth inhibition, apoptosis, and ferroptosis in ILC remains to be explored and will be a component of future studies, as will its role in the additive or synergistic growth suppression achieved by the combination of Riluzole and endocrine therapy. Nevertheless, it is tempting to speculate that agents (like Riluzole) that broadly target rewired cytoskeletal regulatory pathways *and* induce ferroptosis may be particularly effective against ILC.

This study has limitations that are important to consider. First, hormone-responsive ER+ models, particularly of ILC, are limited. In this study, we used three of the four well-established ER+, hormone-responsive ILC lines [71], the ILC-derived HCI-013EI PDX model [15,60], and one of the five PDEs originated from ILC. In all ILC models we tested except for the HCI-013EI PDX, Riluzole plus Fulvestrant provided greater growth inhibition than single-agent Fulvestrant. Expansion of the PDE approach is an important strategy for rapidly diversifying the repertoire of preclinical ILC models [77] to test this and other novel endocrine therapy combinations, essential for a breast cancer subtype that has a significantly greater risk of late recurrence, and worse response to Tamoxifen and the second-generation SERD AZD9496 [12]. Second, Riluzole’s multiple proposed or confirmed mechanisms of action - from inhibition of signaling through GRMs [24] and glutamate release via the glutamate/cystine antiporter SLC7A11 or X_c_^-^ [78], to blockade of voltage-gated sodium channels [79], inhibition of internal ribosome entry site (IRES)-mediated protein synthesis [26], attenuation of RNA polymerase III complex assembly [80], and inhibition of Wnt/β-catenin signaling [81] – present a challenge to readily identifying patients who would benefit most from the drug. Some of this variability is cancer type-specific, with Riluzole action tightly coupled to GRM expression in melanoma [24] and, to some extent, glioblastoma [26], but not in triple-negative breast cancer [27]. While reasonably well-defined in the central nervous system, the interconnected pathways of glutamate release, uptake, and signaling remain understudied in epithelial cells and their pathologies. We posit that future preclinical studies of Riluzole in this context (epithelial tumors generally, and ER+ ILC more specifically) should address these limitations.

## Conclusions

Our data suggest that Riluzole, alone or together with endocrine therapy, may offer therapeutic benefit in some ER+ breast cancers, including ILC, and support optimization and further investigation of Riluzole and its combinations in this setting.

## List of Abbreviations

ER+: estrogen receptor-positive
SERM: selective estrogen receptor modulator
SERD: selective estrogen receptor downregulator
ILC: invasive lobular breast cancer
IDC: invasive ductal breast cancer mGluR
GRM: metabotropic glutamate receptor
ALS: amyotrophic lateral sclerosis
IMEM: improved minimal essential media
FBS: fetal bovine serum
CCS: charcoal-cleared serum
STR: short tandem repeat
DMSO: dimethylsulfoxide
SEM: standard error of the mean
SD: standard deviation
PI: propidium iodide
FITC: fluorescein isothiocyanate
BCA: bicinchoninic acid
PDE: patient-derived explant
PCNA: proliferating cell nuclear antigen
IACUC: institutional animal care and use committee
PDX: patient-derived xenograft
E2: estradiol
SQ: subcutaneous
PO: per os, by mouth
mIHC: multiplex immunohistochemistry
MWT: microwave treatment
AR6: antigen retrieval buffer 6
DAPI: 4’,6-diamidino-2-phenylindole
MDA: malondialdehyde
4-HNE: 4-hydroxynonenal
HSA: highest single agent
ZIP: zero interaction potency
ANOVA: analysis of variance
MM134: MDA-MB-134VI cell line
2D: two-dimensional
NSG: NOD scid gamma
SCID: severe combined immunodeficiency
GPX4: glutathione peroxidase 4
IRES: internal ribosome entry site

## Declarations

### Ethics approval and consent to participate

All animal studies were ethically conducted in accordance with our approved Institutional Animal Care and Use Committee (IACUC) protocols #2018-0005 and #2018-0006. For explant experiments, tissues were collected from discarded surgical samples from UT Southwestern Medical Center (UTSW, Dallas, TX) patients for research purposes after obtaining written informed consent and in accordance with institutional review board-approved protocol (STU-032011–187).

### Consent for publication

Not applicable.

### Availability of data and materials

All data generated or analyzed during this study are included in this published article and its supplementary information files (published in the Zenodo repository, 10.5281/zenodo.7600717).

### Competing interests

The authors declare that they have no competing interests.

### Funding

These studies were supported by the Department of Defense (DoD) Breast Cancer Research Program award W81XWH-17-1-0615 to RBR. Fellowship support for HS was provided by the Tumor Biology Training Grant T32 CA009686 (principal investigator (PI): Dr. Anna T. Riegel). SP received a Georgetown Undergraduate Research Opportunities Program (GUROP) Summer Fellowship. Technical services were provided by the GUMC Animal Models, Flow Cytometry and Cell Sorting, Histopathology and Tissue, and Tissue Culture Shared Resources, which are supported, in part, by NIH/NCI Cancer Center Support Grant P30 CA051008 (PI: Dr. Louis M. Weiner). The content of this article is the sole responsibility of the authors and does not represent the official views of the DoD or NIH.

### Authors’ contributions

AOO and HS contributed to the study design, performed and analyzed experiments, and played a key role in writing the manuscript. SB, SR, and SM contributed to the study design and performed and analyzed experiments. SP performed and analyzed experiments and played a key role in writing the manuscript. YG, MIC, and CB performed and analyzed experiments. AR, HC, and DLB contributed to the study design and performed and analyzed experiments. BMJ and GVR contributed to the study design. RBR contributed to the study design, performed and analyzed experiments, and played a key role in writing the manuscript. All authors read and approved the final manuscript.

## Acknowledgments

The authors would like to thank members of the Riggins laboratory, Drs. Karen Creswell, Michael Johnson, Marc Lippman, Dan Xun (Lombardi Comprehensive Cancer Center, Georgetown University), and Drs. David Lum and Alana Welm (Huntsman Cancer Institute, University of Utah) for sharing reagents, scientific insights, technical assistance, and/or editorial comments on the manuscript.

---

Supplementary Figure Legends, Olukoya et al 2023 (https://doi.org/10.1101/2020.07.30.227561)

**Figure S1.**
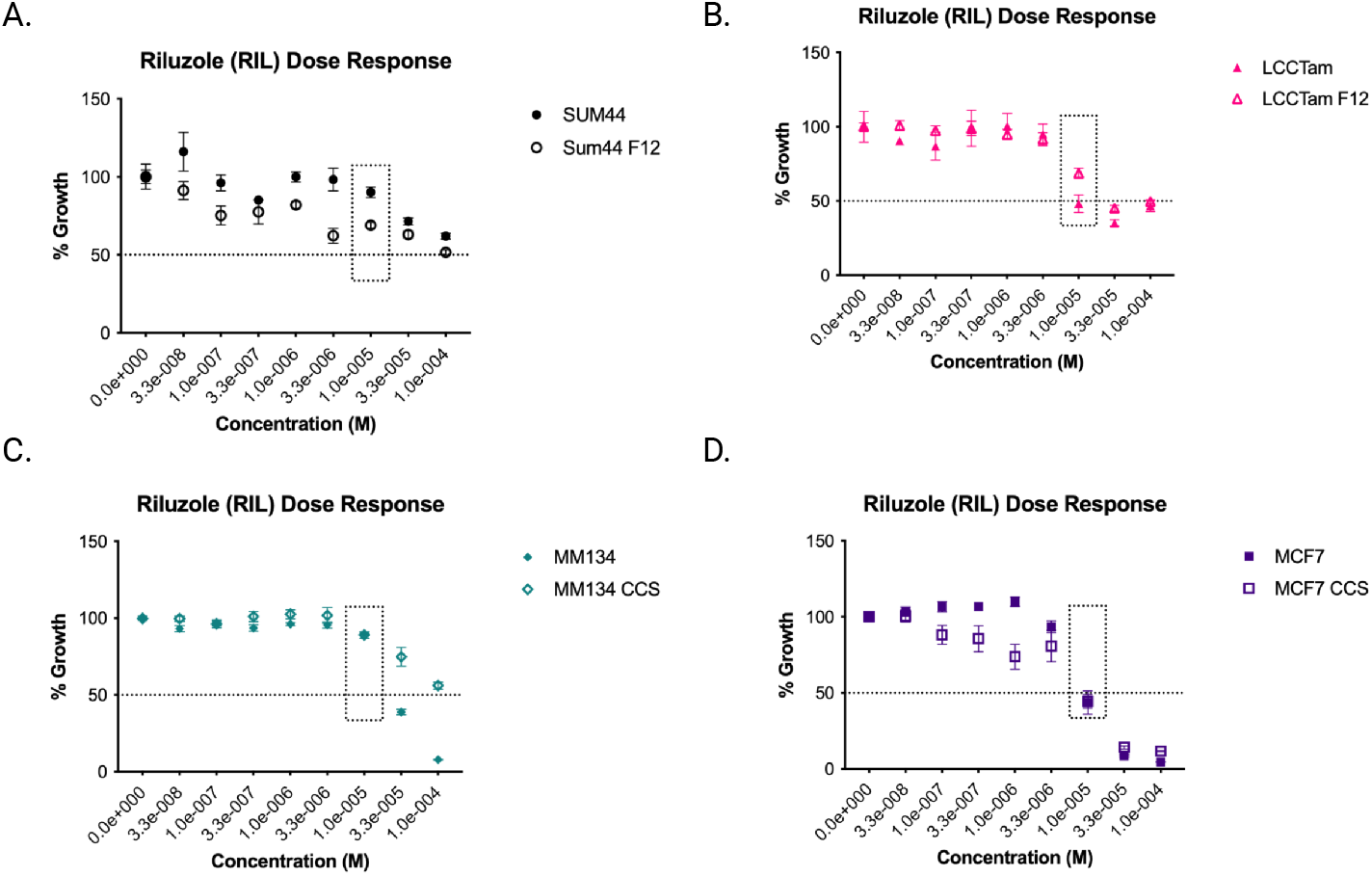
Growth suppression of ER+ breast cancer cell lines by Riluzole in hormone-deprived vs. hormone-replete conditions. In these panels, data already presented in Figure 1A (hormone-replete conditions, filled symbols) were directly compared to hormone-reduced (SUM44 and LCCTam F12 cells, open symbols) or hormone-deprived conditions (MM134 CCS and MCF7 CCS cells, open symbols) as detailed in the Methods section. Cells seeded in 96-well plates were treated with the indicated concentrations of Riluzole (RIL, 33 nM to 100 μM) or DMSO control for 7-8 days prior to staining with crystal violet. Data are presented as mean % growth ± standard error of the mean (SEM) of % growth (vehicle = 100%) for 5-6 technical replicates and represent 2-4 independent biological assays. The dotted line box indicates 10 μM RIL, the concentration used for subsequent *in vitro* assays. Data were analyzed by nonlinear regression ([inhibitor] vs. normalized response), yielding the following IC_50_ [M] estimates: SUM44, 1.27e-4 hormone-replete vs. 5.5e-5 hormone-reduced; LCCTam, 2.13e-5 hormone-replete vs. 3.96e-5 hormone-reduced; MM134, 2.73e-5 hormone-replete vs. 1.16e-4 hormone-deprived; MCF7, 1.09e-5 hormone-replete vs. 7.69e-6 hormone-deprived.

**Figure S2.**
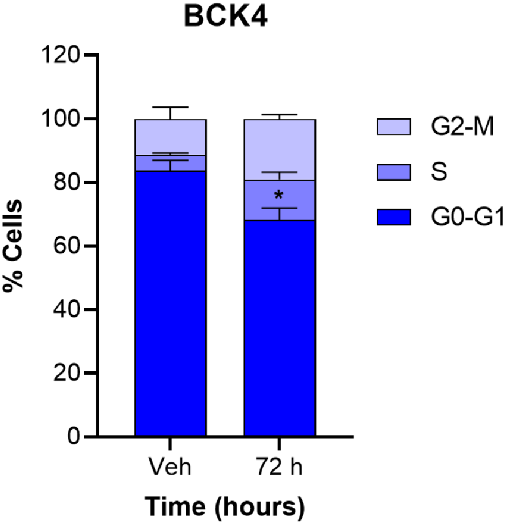
Cell cycle analysis of Riluzole-treated BCK4 cells. Cells seeded in 6-well plates were treated with 10 μM Riluzole or DMSO control (vehicle, Veh) for the indicated times prior to collection, fixation, staining, and cell cycle analysis. Data are presented as mean % cells ± SD for 3-4 independent biological assays and analyzed by two-way ANOVA followed by either Sidak’s (single time point) or Dunnett’s (multiple time points) multiple comparisons tests BCK4: *p=0.044.

**Figure S3.**
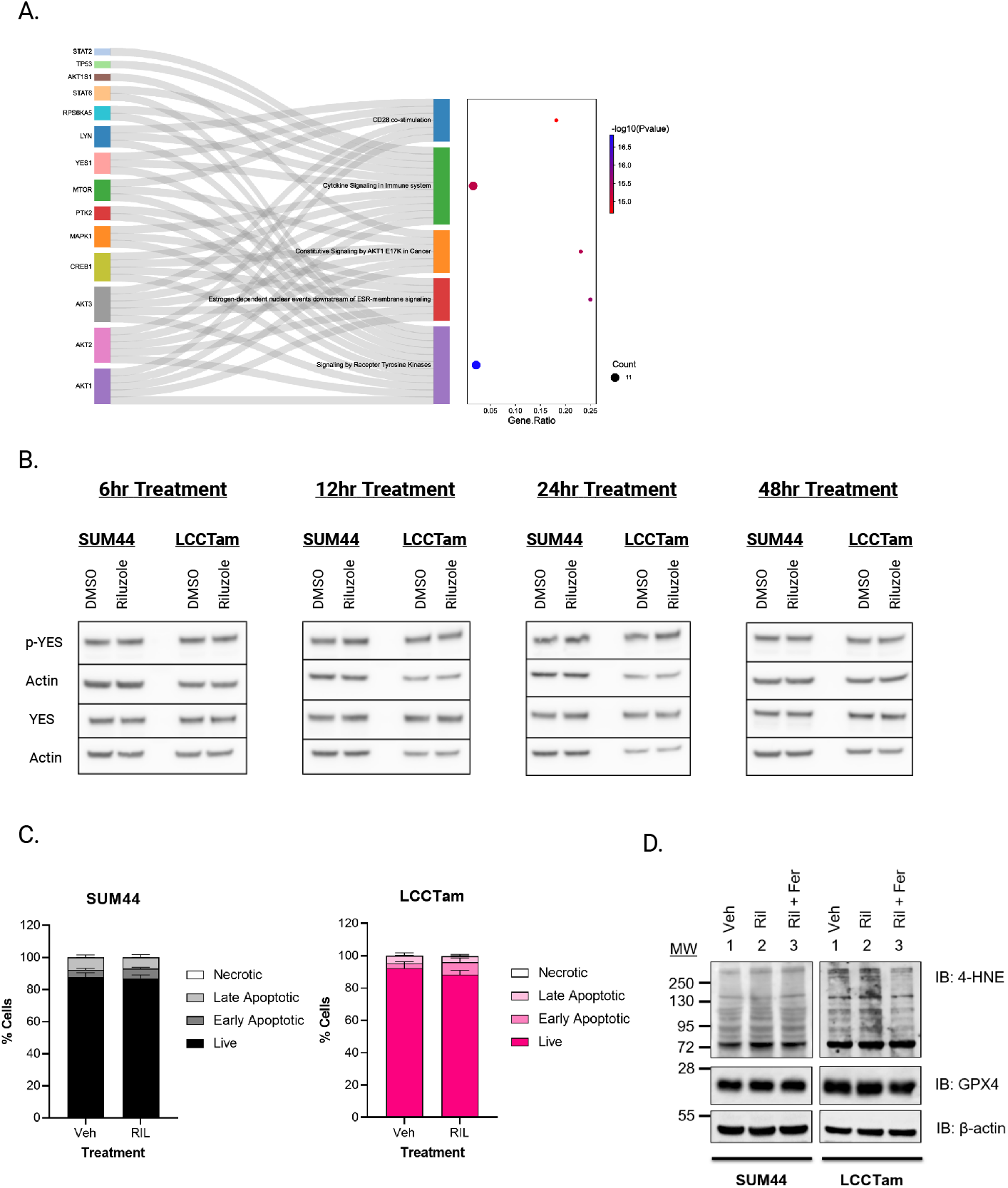
Effect of Riluzole on kinase phosphorylation, cell death, and expression of ferroptosis marker 4-HNE. **A**, SRPlot analysis of gene symbols corresponding to the dephosphorylated kinases and substrates in Riluzole-treated LCCTam cells shown in Figure 3A. False discovery rate (FDR) corrected p values are rank-ordered in -log(10) scale. **B**, Sum44, and LCCTam cells seeded in 6-well plates were treated with 10 μM Riluzole or DMSO control at several time points (6,12, 24, and 48 hours). After which, cells were collected, lysed, and western blot analysis was performed to test for expression and phosphorylation of Yes Y426 phosphorylation. The data are presented as images showing expression levels. **C**, Cells seeded in 6-well plates were treated with 10 μM Riluzole or DMSO control (vehicle, Veh) for 2 days prior to staining with Annexin V and PI. The percent of cells that are live (PI^-^, annexin V^-^), early apoptotic (PI^-^, annexin V^+^), late apoptotic (PI^+^, annexin V^+^), and necrotic (PI^+^, annexin V^-^) are shown. Data are presented as mean % cells ± SD for 3 (SUM44) or 4 (LCCTam) independent biological assays, and analyzed by two-way ANOVA followed by Sidak’s multiple comparisons test. **D**, Sum44, and LCCTam cells seeded in 6-well plates were treated with control (DMSO), Riluzole (10 μM), or a combination of Riluzole and Ferrostatin-1(10 μM) for 48hr. Posttreatment, cells were collected, lysed, and western blot analysis was performed to test for expression of 4-Hydroxynonenal (4HNE). The data are presented as images showing expression levels.

**Figure S4.**
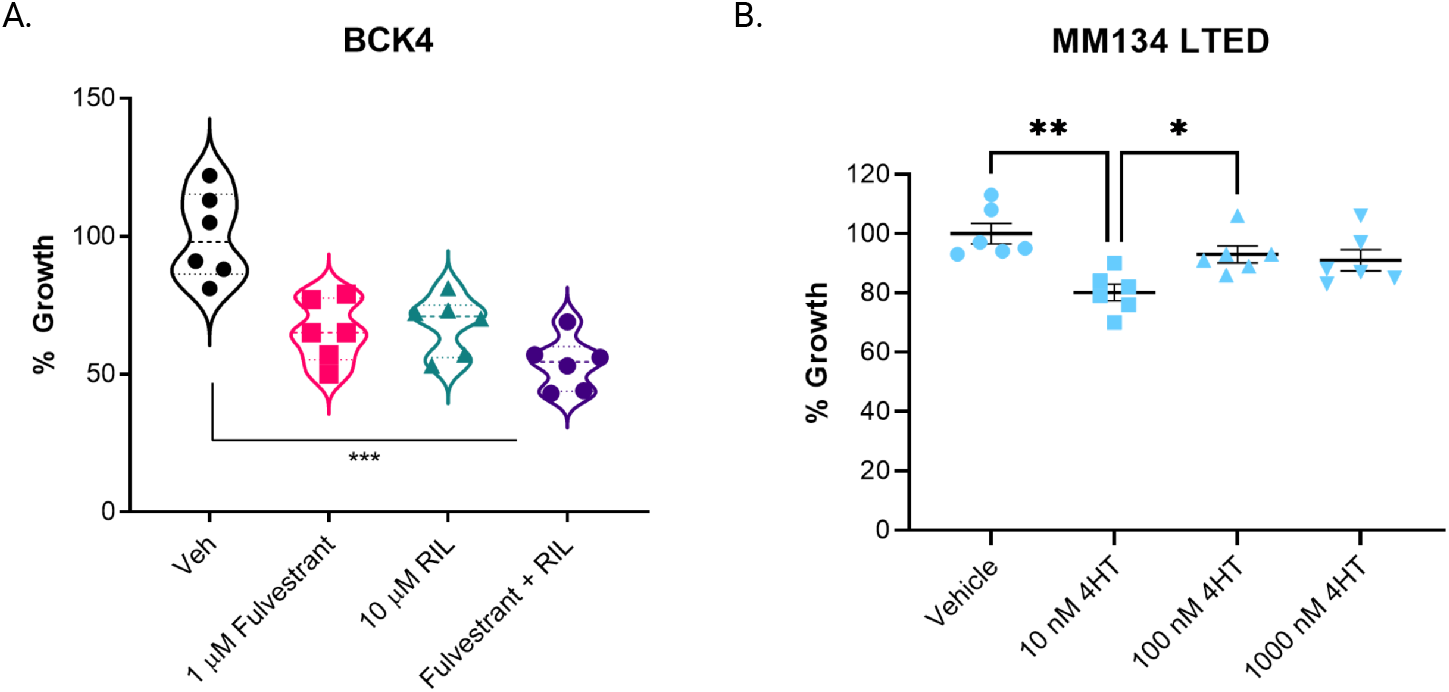
Cell growth assays in BCK4 and MM134 LTED cells. **A**, Cells seeded in 96-well plates were treated with 1 μM Fulvestrant, 10 μM Riluzole (RIL), the combination, or DMSO control (vehicle, Veh) for 7-8 days prior to staining with crystal violet. Data are presented as median with upper/lower quartiles of % growth for 5-6 technical replicates and represent 2-4 independent biological assays. Data were analyzed by the Kruskal-Wallis test, followed by Dunn’s multiple comparison test. **B**, Cells seeded in 96-well plates were treated with 1 μM Fulvestrant, 10 μM Riluzole (RIL), the combination, or DMSO control (vehicle, Veh) for 7-8 days prior to staining with crystal violet. Data are processed as the mean % growth for 5-6 technical replicates and represent 2-4 independent biological assays. The % growth data of a representative single technical replicate is presented in the graph showing the effecting of different concentrations of fulvestrant on MM134 LTED.

**Figure S5.**
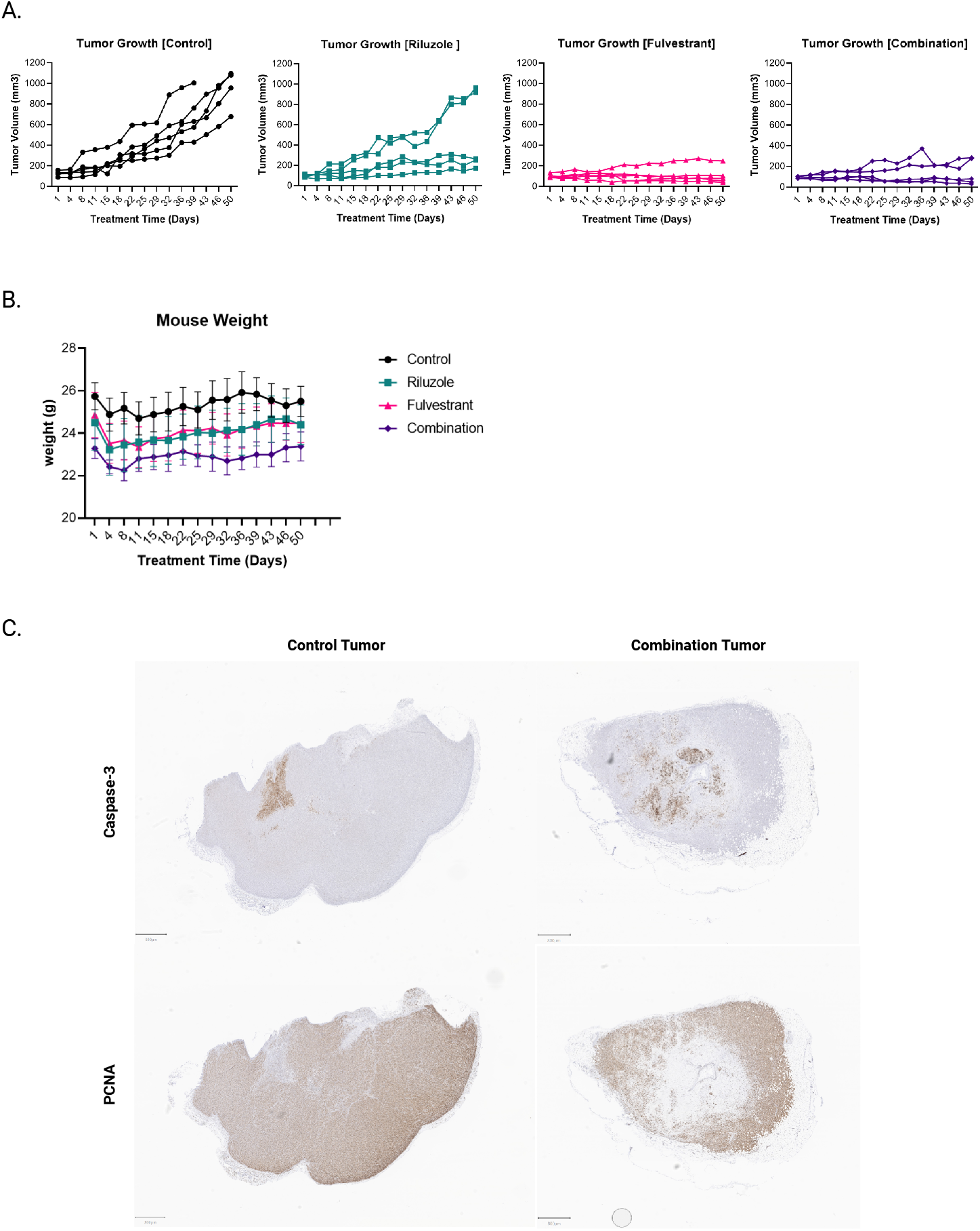
Effect of Riluzole plus Fulvestrant on individual HCI-013EI tumor growth and mouse body weight. **A**, Data already presented in Figure 6A as mean tumor volume ± SEM are re-graphed to show the growth of each individual tumor. **B**, Mouse weights of experimental animals for whom tumor volume measurements are shown in panel A and Figure 6A are presented as mean weight ± SEM for each treatment group. **C**, Representative image of tumor presented in Figure S5A, which were formalin-fixed, paraffin-embedded, sectioned, and stained with proliferating cell nuclear antigen (PCNA) and Caspase-3 by IHC.

**Figure S6.**
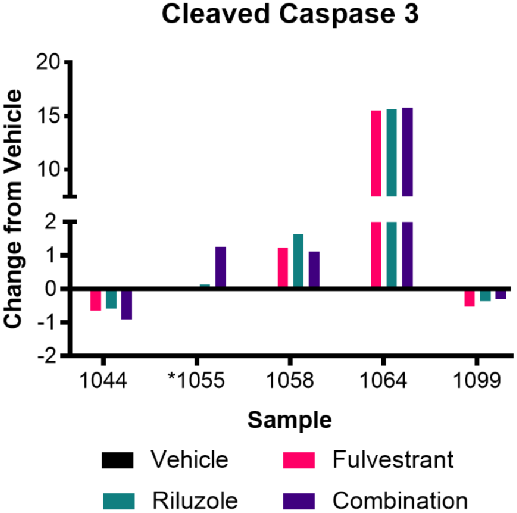
Effect of Riluzole plus Fulvestrant on apoptosis in patient-derived explants (PDEs). PDEs were treated with 100 nM Fulvestrant, 10 μM Riluzole, the combination, or DMSO control (vehicle) for 2 days prior to formalin fixation, paraffin embedding, sectioning, and staining for cleaved caspase 3 by IHC. Data are presented as change relative to vehicle (set to 0) for each explant and analyzed by one-sample t-test vs. 0 (vehicle).

## Notes

Conflicts of Interest: RBR is an Associated Editor for the Journal of the Endocrine Society. All other authors have nothing to disclose related to this Work.

https://doi.org/10.5281/zenodo.7600717

### Summary of Updates

New data and additional authors

## References

1. Siegel RL, Miller KD, Wagle NS, Jemal A. Cancer statistics, 2023. CA Cancer J Clin. 2023;73:17–48.

2. Burstein HJ, Lacchetti C, Anderson H, Buchholz TA, Davidson NE, Gelmon KA, et al. Adjuvant Endocrine Therapy for Women With Hormone Receptor-Positive Breast Cancer: ASCO Clinical Practice Guideline Focused Update. J Clin Oncol. 2019;37:423–38.

3. Hanker AB, Sudhan DR, Arteaga CL. Overcoming Endocrine Resistance in Breast Cancer. Cancer Cell. 2020;37:496–513.

4. Ciriello G, Gatza ML, Beck AH, Wilkerson MD, Rhie SK, Pastore A, et al. Comprehensive Molecular Portraits of Invasive Lobular Breast Cancer. Cell. 2015;163:506–19.

5. Desmedt C, Zoppoli G, Gundem G, Pruneri G, Larsimont D, Fornili M, et al. Genomic Characterization of Primary Invasive Lobular Breast Cancer. J Clin Oncol. 2016;34:1872–81.

6. Michaut M, Chin SF, Majewski I, Severson TM, Bismeijer T, de Koning L, et al. Integration of genomic, transcriptomic and proteomic data identifies two biologically distinct subtypes of invasive lobular breast cancer. Sci Rep. 2016;6:18517.

7. Pestalozzi BC, Zahrieh D, Mallon E, Gusterson BA, Price KN, Gelber RD, et al. Distinct clinical and prognostic features of infiltrating lobular carcinoma of the breast: combined results of 15 International Breast Cancer Study Group clinical trials. J Clin Oncol. 2008;26:3006–14.

8. Adachi Y, Ishiguro J, Kotani H, Hisada T, Ichikawa M, Gondo N, et al. Comparison of clinical outcomes between luminal invasive ductal carcinoma and luminal invasive lobular carcinoma. BMC Cancer. 2016;16:248.

9. Metzger Filho O, Giobbie-Hurder A, Mallon E, Gusterson B, Viale G, Winer EP, et al. Relative Effectiveness of Letrozole Compared With Tamoxifen for Patients With Lobular Carcinoma in the BIG 1-98 Trial. J Clin Oncol. 2015;33:2772–9.

10. Knauer M, Gruber C, Dietze O, Greil R, Stoger H, Rudas M, et al. Abstract S2-06: Survival advantage of anastrozol compared to tamoxifen for lobular breast cancer in the ABCSG-8 study. Cancer Res; 2015. p. Abstract numer S2–06.

11. Strasser-Weippl K, Sudan G, Ramjeesingh R, Shepherd L, et al. Outcomes of invasive ductal (ID) or invasive lobular (IL) early stage breast cancer in women treated with anastrozole or exemestane in the Canadian cancer trials Group MA.27. J Clin Oncol; 2016. p. suppl; abstr 521.

12. Sreekumar S, Levine KM, Sikora MJ, Chen J, Tasdemir N, Carter D, et al. Differential regulation and targeting of estrogen receptor α turnover in invasive lobular breast carcinoma. Endocrinology. 2020;

13. Riggins RB, Lan JP, Zhu Y, Klimach U, Zwart A, Cavalli LR, et al. ERRgamma Mediates Tamoxifen Resistance in Novel Models of Invasive Lobular Breast Cancer. Cancer Res. 2008;68:8908–17.

14. Stires H, Heckler MM, Fu X, Li Z, Grasso CS, Quist MJ, et al. Integrated molecular analysis of Tamoxifen-resistant invasive lobular breast cancer cells identifies MAPK and GRM/mGluR signaling as therapeutic vulnerabilities. Mol Cell Endocrinol. 2017/09/25 ed. 2018;471:105–17.

15. Sikora MJ, Cooper KL, Bahreini A, Luthra S, Wang G, Chandran UR, et al. Invasive lobular carcinoma cell lines are characterized by unique estrogen-mediated gene expression patterns and altered tamoxifen response. Cancer Res. 2014/01/14 ed. 2014;74:1463–74.

16. Sikora MJ, Jacobsen BM, Levine K, Chen J, Davidson NE, Lee AV, et al. WNT4 mediates estrogen receptor signaling and endocrine resistance in invasive lobular carcinoma cell lines. Breast Cancer Res. 2016;18:92.

17. Sokol ES, Feng YX, Jin DX, Basudan A, Lee AV, Atkinson JM, et al. Loss of function of NF1 is a mechanism of acquired resistance to endocrine therapy in lobular breast cancer. Ann Oncol. 2019;30:115–23.

18. Bossart EA, Tasdemir N, Sikora MJ, Bahreini A, Levine KM, Chen J, et al. SNAIL is induced by tamoxifen and leads to growth inhibition in invasive lobular breast carcinoma. Breast Cancer Res Treat. 2019;175:327–37.

19. Levine KM, Priedigkeit N, Basudan A, Tasdemir N, Sikora MJ, Sokol ES, et al. FGFR4 overexpression and hotspot mutations in metastatic ER+ breast cancer are enriched in the lobular subtype. NPJ Breast Cancer. 2019;5:19.

20. Du T, Sikora MJ, Levine KM, Tasdemir N, Riggins RB, Wendell SG, et al. Key regulators of lipid metabolism drive endocrine resistance in invasive lobular breast cancer. Breast Cancer Res. 2018;20:106.

21. Du T, Zhu L, Levine KM, Tasdemir N, Lee AV, Vignali DAA, et al. Invasive lobular and ductal breast carcinoma differ in immune response, protein translation efficiency and metabolism. Sci Rep. 2018;8:7205.

22. Ulaner GA, Goldman DA, Gönen M, Pham H, Castillo R, Lyashchenko SK, et al. Initial Results of a Prospective Clinical Trial of 18F-Fluciclovine PET/CT in Newly Diagnosed Invasive Ductal and Invasive Lobular Breast Cancers. J Nucl Med. 2016;57:1350–6.

23. Ulaner GA, Goldman DA, Corben A, Lyashchenko SK, Gönen M, Lewis JS, et al. Prospective Clinical Trial of 18F-Fluciclovine PET/CT for Determining the Response to Neoadjuvant Therapy in Invasive Ductal and Invasive Lobular Breast Cancers. J Nucl Med. 2017;58:1037–42.

24. Namkoong J, Shin S-S, Lee HJ, Marín YE, Wall BA, Goydos JS, et al. Metabotropic glutamate receptor 1 and glutamate signaling in human melanoma. Cancer Res. 2007;67:2298–305.

25. Khan AJ, Wall B, Ahlawat S, Green C, Schiff D, Mehnert JM, et al. Riluzole enhances ionizing radiation-induced cytotoxicity in human melanoma cells that ectopically express metabotropic glutamate receptor 1 in vitro and in vivo. Clin Cancer Res. 2011;17:1807–14.

26. Benavides-Serrato A, Saunders JT, Holmes B, Nishimura RN, Lichtenstein A, Gera J. Repurposing Potential of Riluzole as an ITAF Inhibitor in mTOR Therapy Resistant Glioblastoma. Int J Mol Sci. 2020;21.

27. Speyer CL, Nassar MA, Hachem AH, Bukhsh MA, Jafry WS, Khansa RM, et al. Riluzole mediates anti-tumor properties in breast cancer cells independent of metabotropic glutamate receptor-1. Breast Cancer Res Treat. 2016;157:217–28.

28. Dolfi SC, Medina DJ, Kareddula A, Paratala B, Rose A, Dhami J, et al. Riluzole exerts distinct antitumor effects from a metabotropic glutamate receptor 1-specific inhibitor on breast cancer cells. Oncotarget. 2017;8:44639–53.

29. Banda M, Speyer CL, Semma SN, Osuala KO, Kounalakis N, Torres Torres KE, et al. Metabotropic glutamate receptor-1 contributes to progression in triple negative breast cancer. PLoS One. 2014;9:e81126.

30. Speyer CL, Bukhsh MA, Jafry WS, Sexton RE, Bandyopadhyay S, Gorski DH. Riluzole synergizes with paclitaxel to inhibit cell growth and induce apoptosis in triple-negative breast cancer. Breast Cancer Res Treat. 2017;166:407–19.

31. Teh JL, Shah R, La Cava S, Dolfi SC, Mehta MS, Kongara S, et al. Metabotropic glutamate receptor 1 disrupts mammary acinar architecture and initiates malignant transformation of mammary epithelial cells. Breast Cancer Res Treat. 2015/04/10 ed. 2015;151:57–73.

32. Jambal P, Badtke MM, Harrell JC, Borges VF, Post MD, Sollender GE, et al. Estrogen switches pure mucinous breast cancer to invasive lobular carcinoma with mucinous features. Breast Cancer Res Treat. 2013;137:431–48.

33. Heckler MM, Zeleke TZ, Divekar SD, Fernandez AI, Tiek DM, Woodrick J, et al. Antimitotic activity of DY131 and the estrogen-related receptor beta 2 (ERRβ2) splice variant in breast cancer. Oncotarget. 2016;7:47201–20.

34. Tiek DM, Khatib SA, Trepicchio CJ, Heckler MM, Divekar SD, Sarkaria JN, et al. Estrogen-related receptor β activation and isoform shifting by cdc2-like kinase inhibition restricts migration and intracranial tumor growth in glioblastoma. FASEB J. 2019;33:13476–91.

35. Heckler MM, Riggins RB. ERRβ splice variants differentially regulate cell cycle progression. Cell Cycle. 2015;14:31–45.

36. Schindelin J, Arganda-Carreras I, Frise E, Kaynig V, Longair M, Pietzsch T, et al. Fiji: an open-source platform for biological-image analysis. Nature Methods. Nature Publishing Group; 2012;9:676–82.

37. Bankhead P, Loughrey MB, Fernández JA, Dombrowski Y, McArt DG, Dunne PD, et al. QuPath: Open source software for digital pathology image analysis. Sci Rep. Nature Publishing Group; 2017;7:16878.

38. Ranjit S, Malacrida L, Jameson DM, Gratton E. Fit-free analysis of fluorescence lifetime imaging data using the phasor approach. Nat Protoc. 2018/09/08 ed. 2018;13:1979–2004.

39. Dvornikov A, Malacrida L, Gratton E. The DIVER Microscope for Imaging in Scattering Media. Methods Protoc [Internet]. 2019/06/27 ed. 2019;2. Available from: https://www.ncbi.nlm.nih.gov/pubmed/31234383

40. Ranjit S, Lanzano L, Gratton E. Mapping diffusion in a living cell via the phasor approach. Biophys J. 2014;107:2775–85.

41. Ranjit S, Dvornikov A, Dobrinskikh E, Wang X, Luo Y, Levi M, et al. Measuring the effect of a Western diet on liver tissue architecture by FLIM autofluorescence and harmonic generation microscopy. Biomed Opt Express. 2017;8:3143–54.

42. Hedde PN, Ranjit S, Gratton E. 3D fluorescence anisotropy imaging using selective plane illumination microscopy. Opt Express. 2015;23:22308–17.

43. Malacrida L, Ranjit S, Jameson DM, Gratton E. The Phasor Plot: A Universal Circle to Advance Fluorescence Lifetime Analysis and Interpretation. Annu Rev Biophys. 2021;50:575–93.

44. Digman MA, Caiolfa VR, Zamai M, Gratton E. The phasor approach to fluorescence lifetime imaging analysis. Biophys J. 2008;94:L14–16.

45. Clayton AHA, Hanley QS, Verveer PJ. Graphical representation and multicomponent analysis of single-frequency fluorescence lifetime imaging microscopy data. J Microsc. 2004;213:1–5.

46. Redford GI, Clegg RM. Polar plot representation for frequency-domain analysis of fluorescence lifetimes. J Fluoresc. 2005;15:805–15.

47. Stringari C, Edwards RA, Pate KT, Waterman ML, Donovan PJ, Gratton E. Metabolic trajectory of cellular differentiation in small intestine by Phasor Fluorescence Lifetime Microscopy of NADH. Sci Rep. 2012;2:568.

48. Ranjit S, Malacrida L, Stakic M, Gratton E. Determination of the metabolic index using the fluorescence lifetime of free and bound nicotinamide adenine dinucleotide using the phasor approach. J Biophotonics. 2019;12:e201900156.

49. Liggett JR, Kang J, Ranjit S, Rodriguez O, Loh K, Patil D, et al. Oral N-acetylcysteine decreases IFN-γ production and ameliorates ischemia-reperfusion injury in steatotic livers. Front Immunol. 2022;13:898799.

50. Centenera MM, Hickey TE, Jindal S, Ryan NK, Ravindranathan P, Mohammed H, et al. A patient-derived explant (PDE) model of hormone-dependent cancer. Mol Oncol. 2018/08/18 ed. 2018;12:1608–22.

51. Ianevski A, Giri AK, Aittokallio T. SynergyFinder 2.0: visual analytics of multi-drug combination synergies. Nucleic Acids Res. 2020;

52. Yadav B, Wennerberg K, Aittokallio T, Tang J. Searching for Drug Synergy in Complex Dose-Response Landscapes Using an Interaction Potency Model. Comput Struct Biotechnol J. 2015;13:504–13.

53. RECIST 1.1-Update and clarification: From the RECIST committee - PubMed [Internet]. [cited 2023 Jan 12]. Available from: https://pubmed.ncbi.nlm.nih.gov/27189322/

54. Brünner N, Boysen B, Jirus S, Skaar TC, Holst-Hansen C, Lippman J, et al. MCF7/LCC9: an antiestrogen resistant MCF-7 variant in which acquired resistance to the steroidal antiestrogen ICI 182,780 confers an early crossresistance to the non-steroidal antiestrogen tamoxifen. Cancer Research. 1997;57:3486–93.

55. Olukoya AO. Data from: Riluzole suppresses growth and enhances response to endocrine therapy in ER+ breast cancer [Internet]. 2023. Available from: 10.5281/zenodo.7600717

56. Wasielewski M, Elstrodt F, Klijn JG, Berns EM, Schutte M. Thirteen new p53 gene mutants identified among 41 human breast cancer cell lines. Breast Cancer ResTreat. 2006;99:97–101.

57. Rosenberg SA, Niglio SA, Salehomoum N, Chan JL-K, Jeong B-S, Wen Y, et al. Targeting Glutamatergic Signaling and the PI3 Kinase Pathway to Halt Melanoma Progression. Transl Oncol. 2015;8:1–9.

58. Yi J, Zhu J, Wu J, Thompson CB, Jiang X. Oncogenic activation of PI3K-AKT-mTOR signaling suppresses ferroptosis via SREBP-mediated lipogenesis. Proc Natl Acad Sci U S A. 2020;117:31189–97.

59. Liu S, Shi J, Wang L, Huang Y, Zhao B, Ding H, et al. Loss of EMP1 promotes the metastasis of human bladder cancer cells by promoting migration and conferring resistance to ferroptosis through activation of PPAR gamma signaling. Free Radical Biology and Medicine. 2022;189:42–57.

60. Scherer SD, Riggio AI, Haroun F, DeRose YS, Ekiz HA, Fujita M, et al. An immune-humanized patient-derived xenograft model of estrogen-independent, hormone receptor positive metastatic breast cancer. Breast Cancer Res. 2021;23:100.

61. Bahreini A, Li Z, Wang P, Levine KM, Tasdemir N, Cao L, et al. Mutation site and context dependent effects of ESR1 mutation in genome-edited breast cancer cell models. Breast Cancer Research. 2017;19:60.

62. Demas DM, Demo S, Fallah Y, Clarke R, Nephew KP, Althouse S, et al. Glutamine Metabolism Drives Growth in Advanced Hormone Receptor Positive Breast Cancer. Front Oncol. 2019;9:686.

63. Bacci M, Lorito N, Ippolito L, Ramazzotti M, Luti S, Romagnoli S, et al. Reprogramming of Amino Acid Transporters to Support Aspartate and Glutamate Dependency Sustains Endocrine Resistance in Breast Cancer. Cell Rep. 2019/07/04 ed. 2019;28:104–118 e8.

64. Ranjit S, Datta R, Dvornikov A, Gratton E. Multicomponent Analysis of Phasor Plot in a Single Pixel to Calculate Changes of Metabolic Trajectory in Biological Systems. J Phys Chem A. 2019/10/23 ed. 2019;123:9865–73.

65. Shah R, Singh SJ, Eddy K, Filipp FV, Chen S. Concurrent Targeting of Glutaminolysis and Metabotropic Glutamate Receptor 1 (GRM1) Reduces Glutamate Bioavailability in GRM1+ Melanoma. Cancer Res. American Association for Cancer Research; 2019;79:1799–809.

66. Mehnert JM, Silk AW, Lee JH, Dudek L, Jeong B-S, Li J, et al. A phase II trial of riluzole, an antagonist of metabotropic glutamate receptor 1 (GRM1) signaling, in patients with advanced melanoma. Pigment Cell Melanoma Res. 2018;31:534–40.

67. Pelletier JC, Chen S, Bian H, Shah R, Smith GR, Wrobel JE, et al. Dipeptide Prodrugs of the Glutamate Modulator Riluzole. ACS Med Chem Lett. 2018;9:752–6.

68. Silk AW, Saraiya B, Groisberg R, Chan N, Spencer K, Girda E, et al. A phase Ib dose-escalation study of troriluzole (BHV-4157), an oral glutamatergic signaling modulator, in combination with nivolumab in patients with advanced solid tumors. Eur J Med Res. 2022;27:107.

69. Hiscox S, Barnfather P, Hayes E, Bramble P, Christensen J, Nicholson RI, et al. Inhibition of focal adhesion kinase suppresses the adverse phenotype of endocrine-resistant breast cancer cells and improves endocrine response in endocrine-sensitive cells. Breast Cancer Res Treat. 2011;125:659–69.

70. Schwarz LJ, Fox EM, Balko JM, Garrett JT, Kuba MG, Estrada MV, et al. LYN-activating mutations mediate antiestrogen resistance in estrogen receptor-positive breast cancer. J Clin Invest. 2014;124:5490–502.

71. Tasdemir N, Bossart EA, Li Z, Zhu L, Sikora MJ, Levine KM, et al. Comprehensive Phenotypic Characterization of Human Invasive Lobular Carcinoma Cell Lines in 2D and 3D Cultures. Cancer Res. 2018;78:6209–22.

72. Bruner HC, Derksen PWB. Loss of E-Cadherin-Dependent Cell-Cell Adhesion and the Development and Progression of Cancer. Cold Spring Harb Perspect Biol. 2018;10.

73. Paoli P, Giannoni E, Chiarugi P. Anoikis molecular pathways and its role in cancer progression. Biochimica et Biophysica Acta (BBA) - Molecular Cell Research. 2013;1833:3481–98.

74. Zou D, Yoon H-S, Anjomshoaa A, Perez D, Fukuzawa R, Guilford P, et al. Increased levels of active c-Src distinguish invasive from in situ lobular lesions. Breast Cancer Res. 2009;11:R45.

75. Panzilius E, Holstein F, Dehairs J, Planque M, von Toerne C, Koenig A-C, et al. Cell density-dependent ferroptosis in breast cancer is induced by accumulation of polyunsaturated fatty acid-enriched triacylglycerides. bioRxiv. 2019;417949.

76. Wu J, Minikes AM, Gao M, Bian H, Li Y, Stockwell BR, et al. Intercellular interaction dictates cancer cell ferroptosis via NF2-YAP signalling. Nature. 2019/07/26 ed. 2019;572:402–6.

77. Bahnassy S, Sikora MJ, Riggins RB. Unlocking the Mysteries of Lobular Breast Cancer Biology Needs the Right Combination of Preclinical Models. Mol Cancer Res. 2022;20:837–40.

78. Shin S-S, Jeong B-S, Wall BA, Li J, Shan NL, Wen Y, et al. Participation of xCT in melanoma cell proliferation in vitro and tumorigenesis in vivo. Oncogenesis. 2018;7:86.

79. Djamgoz MBA, Onkal R. Persistent current blockers of voltage-gated sodium channels: a clinical opportunity for controlling metastatic disease. Recent Pat Anticancer Drug Discov. 2013;8:66–84.

80. Pinard M, Dastpeyman S, Poitras C, Bernard G, Gauthier M-S, Coulombe B. Riluzole partially restores RNA polymerase III complex assembly in cells expressing the leukodystrophy-causative variant POLR3B R103H. Mol Brain. 2022;15:98.

81. Roy SK, Ma Y, Lam BQ, Shrivastava A, Srivastav S, Shankar S, et al. Riluzole regulates pancreatic cancer cell metabolism by suppressing the Wnt-β-catenin pathway. Sci Rep. 2022;12:11062.

